# Evaluating single-cell variability in proteasomal decay

**DOI:** 10.1101/2023.08.22.554358

**Authors:** Sukanya Das, Abhyudai Singh, Premal Shah

## Abstract

Gene expression is a stochastic process that leads to variability in mRNA and protein abundances even within an isogenic population of cells grown in the same environment. This variation, often called gene-expression noise, has typically been attributed to transcriptional and translational processes while ignoring the contributions of protein decay variability across cells. Here we estimate the single-cell protein decay rates of two degron GFPs in *Saccharomyces cerevisiae* using time-lapse microscopy. We find substantial cell-to-cell variability in the decay rates of the degron GFPs. We evaluate cellular features that explain the variability in the proteasomal decay and find that the amount of 20s catalytic beta subunit of the proteasome marginally explains the observed variability in the degron GFP half-lives. We propose alternate hypotheses that might explain the observed variability in the decay of the two degron GFPs. Overall, our study highlights the importance of studying the kinetics of the decay process at single-cell resolution and that decay rates vary at the single-cell level, and that the decay process is stochastic. A complex model of decay dynamics must be included when modeling stochastic gene expression to estimate gene expression noise.

## Introduction

Gene expression is a cascade of biochemical reactions involving transcription, translation, and degradation of mRNAs and proteins. Each of these processes is inherently stochastic. The stochastic nature leads to variability in the number of mRNA and protein molecules even in an isogenic population of cells grown in the same environment, often called gene expression noise [1–7].

Gene expression noise partly arises due to features inherent to a gene, such as its promoter sequences [8–15], chromosomal architecture [16–18], genomic location [19–21], the secondary structure of mRNA [22–24], and the protein encoded [25], typically referred to as intrinsic sources [26] and noise due to these sources are referred to as intrinsic noise in gene expression. In addition to gene-specific sources of variation, gene expression noise can arise due to global cellular features such as the cell cycle stage [27–30], cellular age [31], asymmetric partitioning of cellular components during cell division [32], differences in the number of molecules governing gene expression, like the amount of transcription factors[33–35], polymerases, ribosomes, tRNAs[36], mRNA decay machinery [37] and the number of proteasomes[38], in a cell. These sources can influence the dynamics of the reactions involved in gene expression, and noise originating from these sources is called extrinsic noise. The dynamics of transcription [6, 17, 19, 39–42] and translation [7, 22, 43] are thought to be the predominant source of the noise in gene expression. For instance, the discontinuous production of mRNAs, known as transcriptional bursts, due to periods of active and inactive transcription, is widely considered to modulate gene expression noise [39, 40, 42, 44–46].

While the kinetics of transcription and translation lead to gene expression noise by influencing the production of mRNA and protein, degradation of these molecules can also impact gene expression noise due to the dynamic nature of mRNA and protein degradation [47–50]. Furthermore, many theoretical models of stochastic gene expression consider protein degradation to be slow relative to the birth and death of mRNA, simplifying the model by assuming that the protein half-lives are in the order of hours [51, 52]. While this might be true for most proteins, a non-trivial proportion of the *S. cerevesiae* proteome degrades in the order of minutes [53, 54]. The amount of these fast-decaying proteins was shown to be regulated by protein degradation and not by transcription or translation [53]. As a result, the previous studies on gene expression noise might have underestimated the contribution of protein decay while overestimating the contribution of transcription and translation processes to noise. Additionally, one theoretical model for the decomposition of noise in gene expression has shown that the process of degradation is responsible for up to 20-40% of the observed variance [55]. Regulated protein degradation can also attenuate noise in the expression of heat shock chaperones responding to high-temperature stress in *E. coli* [56]. These theoretical studies point toward the potential role of protein degradation in modulating noise, but there is a lack of experimental work assessing this role. It was recently shown that low noise levels in the machinery involved in the Nonsense-mediated mRNA decay reduced the noise in short-lived mRNAs [37]. While an important contribution to how the decay process can affect global gene expression noise, the study does not address the single-cell variation in mRNA decay dynamics. Another recent study found substantial cell-to-cell variation in proteasomal decay rates, which was cell-cycle dependent in mammalian cells [38].

Experimental studies measuring single-cell abundances of mRNA transcripts and proteins do so at a single snapshot of time, lacking the temporal resolution needed to study the kinetics of processes at the single-cell resolution. Most studies addressing stochastic gene expression estimate a single rate constant for a given process using quantitative measurements from various time points, inferring a single rate constant for a population of cells. These studies then change the rates of processes by introducing either a perturbation[17, 57] or by changing the concentration of an inducer. The output of these perturbations is reflected in a change in the number of mRNA and protein molecules in single cells in turn affecting the single-cell variability in these molecules. The role of the perturbed process on cellular heterogeneity is then inferred from these static measurements. What we don’t know is how the kinetics of the processes of gene expression vary at the single-cell level. Does the rate constant of a process itself vary from one cell to another? Furthermore, if the rates do vary, what are some of the cellular features that can explain the observed variability?

To address these questions, we studied the single-cell kinetics of the decay of two degron GFPs using time-lapse fluorescent microscopy. Using the time-series GFP trace of single cells, we estimated the single-cell rate of proteasomal decay of two degron GFPs each getting targeted to the proteasome in either ubiquitin-independent or dependent manner. We found substantial cell-to-cell variability in the protein decay rate, and the degree of variation was independent of the pathway to the proteasome. We then evaluated various cellular features that might explain the observed variation in single-cell decay rates. The initial amount of GFP in a cell was the biggest predictor of the protein decay rate in a cell. Interestingly, we find that the amount of a catalytic proteasomal subunit did not correlate with the single-cell decay rates. Overall, our study quantifies the extent of variability in the proteasomal decay in yeast and helps shed light on its potential sources.

## Results

### Estimating single cell proteasomal decay rates

To study the cell-to-cell variability in protein decay rates, we used two destabilized GFPs. One destabilized GFP named yeGFP-mODC contained a mouse Ornithine decarboxylase (mODC) PEST sequence [58] fused to the C-terminal of the yeast-enhanced GFP (yeGFP). This targets the GFP to the proteasome in a ubiquitin-independent manner [58] and destabilizes the protein in yeast [59]. The other destabilized GFP used in the study has the PEST sequence from the yeast protein Cyclin-2 fused to the c-terminal of yeGFP [60, 61]. This targets the protein for proteasomal degradation via the ubiquitin pathway [62]. The fusion of this pest sequence to the C-terminal of yeGFP was previously shown to reduce the half-life of yeGFP from 7 hrs to 30 mins [63]. The GFP expression cassettes were genomically integrated at the *LEU2* locus and transcriptionally expressed by a galactose inducible *GAL1* promoter. To decouple the effects of transcription and translation from the protein decay, transcription of new GFP mRNAs was inhibited by glucose, and translation was stalled by cycloheximide (CHX), preventing the production of new GFP molecules. Single cells were segmented and tracked, and the mean GFP intensity trace for each cell was quantified (**Figure 1A, Figure 2B**). Various filtering criteria were instituted to remove dead cells, blurry cells, and cells with GFP intensity near autofluorescence control intensities (*see materials and methods/Microscopy*). The cells which passed the filtering criteria were used to estimate the single-cell GFP decay rates. The bulk population-level degradation kinetics of yeGFP-mODC and yeGFP-CLN2 is shown in (**Figure 2A**). The raw data GFP intensity (background subtracted) in the cells expressing the degron GFPs against the autofluorescence of the parental strain is shown in **Supplemental figure S1**. The reduction in the mean GFP intensity of the population of cells is shown in **Supplemental figure S2**. The single-cell GFP traces of three cells with varying GFP decays are shown in (**Figure 2B**).

**Figure 1.**
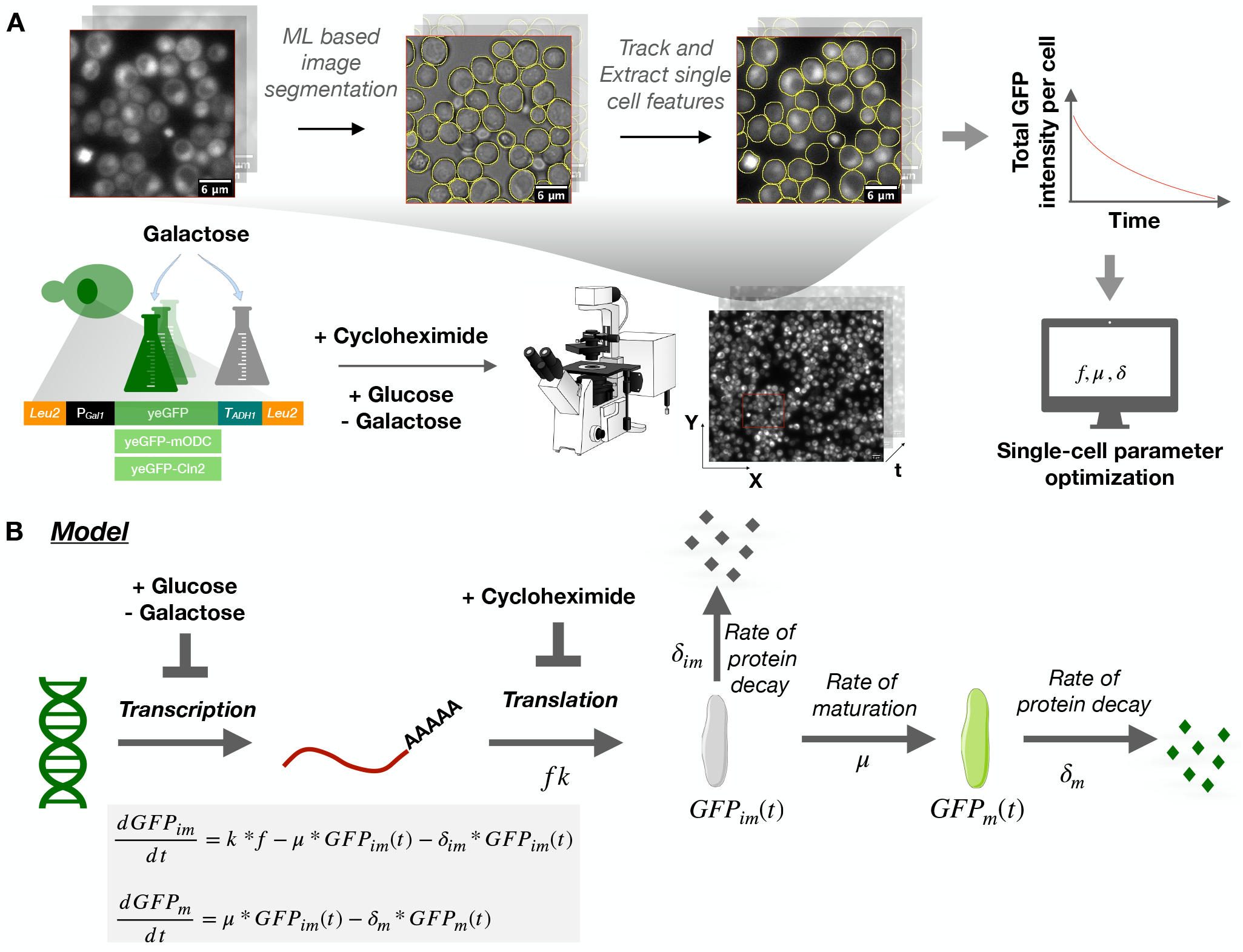
Experimental method to estimate the single-cell rate of protein degradation. **A:** Experimental framework to estimate the single cell degron GFP half-lives. Three different GFP species with varying decay rates and mechanisms are studied. After 3hrs of induction of GFP, transcription is inhibited by changing the carbon source from galactose to glucose, and the translation is inhibited using cycloheximide (CHX). Cells were immediately imaged using a fluorescent microscope. Single cells were segmented using the YeastSpotter tool [96], tracked in every image from the time-lapse, and single-cell features like area, fluorescent intensities, cellular shape, etc. were extracted. The single-cell GFP intensity trace was used to estimate the single-cell rates of proteasomal degradation. **B**: The mechanistic model of GFP decay. Upon shutdown of transcription (+Glucose) and translation (+CHX), we assume a low amount of new GFP synthesis by leaky translation at the rate of *fk* due to the reversible nature of CHX. We assume that immature GFP molecules [*GFP*_*im*_(*t*)] undergo maturation at the rate of *µ* and add to the pool of mature GFP *GFP*_*m*_(*t*) in the cell. The proteasome decays immature and mature GFP molecules at the rate of *δ*_*im*_ and *δ*_*m*_, respectively. The rate of change of the two GFP molecules can be written as the ordinary differential equation shown. We solve these equations assuming that immature and mature GFP decay at the same rate, and with the initial conditions shown in **eq 3-4**. See Materials and Methods and Model fitting for complete details.

**Figure 2.**
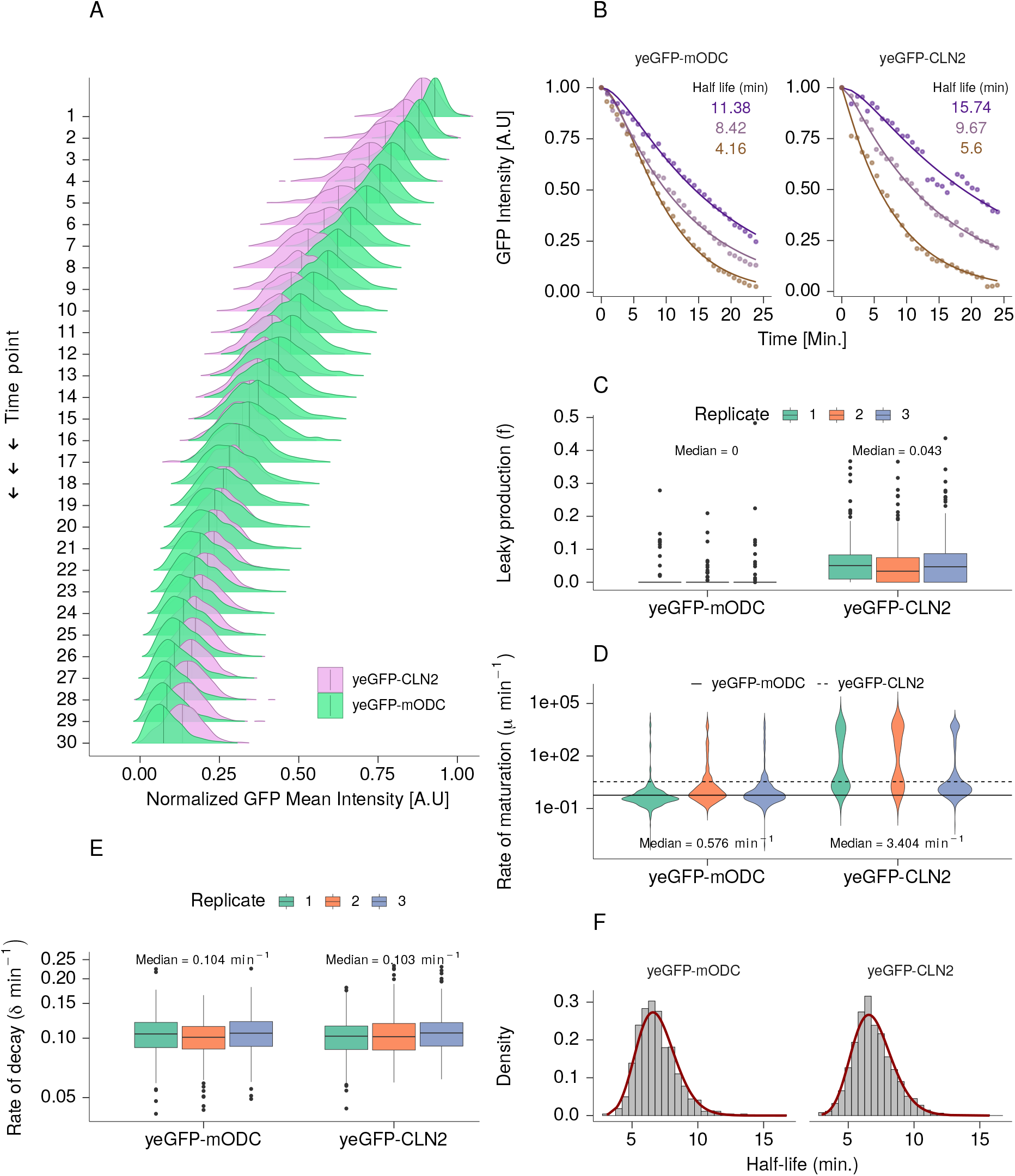
Estimation of single cell degron GFP Half-lives. **A:** Time series of GFP intensities. For each time point, the GFP intensities of single cells are normalized to the initial GFP intensity. Each image was taken approximately at 40-50 sec intervals. Vertical lines represent the median normalized GFP intensity at each time point. **B:** GFP decay in single-cells. The GFP intensity trace for three random cells with varying GFP half-lives is plotted. The experimental data is shown as points, and the model fit the corresponding cell is shown as solid lines. **C:** Box plots of the leaky production of GFP (*f*). The *f* parameter values for single cells are estimated from the mechanistic model of decay eq.5. The leaky production of GFP (*f*) corresponds to the fraction of translation occurring without translation inhibition. The median values are shown. The whiskers extend to 1.5 times the interquartile range. The median *f* value of the pooled data for each degron GFP is shown. **D:** Rate of maturation (*µ min*^*−*1^) estimated from eq.5. The single-cell values of maturation rate are shown as violin plots with a median GFP maturation rate of 0.58 *min*^*−*1^ for yeGFP-mODC and 3.4 *min*^*−*1^ for yeGFP-CLN2. **E:** Rate of GFP decay (*δ min*^*−*1^). The single-cell values of the degron GFP rate of decay estimated from eq.5 are represented as box plots. The three colors in panels C-E correspond to three different biological replicates. The three biological replicates are very similar to each other. Hence they were pooled for the remaining analysis. **F**: Distribution of single-cell degron GFP half-lives. The half-lives were calculated as 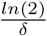 and were fitted to a gamma distribution (red line).

To estimate the single-cell GFP decay rates and thus the half-lives, we developed a mechanistic model of GFP decay (**Figure 1B**, *see materials and methods/Model*). Initially, galactose induces the transcription of the GFP mRNAs, which are translated to proteins. Let *k* be the rate at which new GFP proteins are produced at steady-state. To isolate the GFP decay dynamics from synthesis, the production of new GFP proteins was inhibited by adding glucose, a potent transcription inhibitor of the galactose promoter. In addition, we added cycloheximide (CHX), a reversible translation elongation inhibitor to block the synthesis of new GFP proteins from existing mRNAs. Throughout the timelapse, a residual translation of GFP might occur at a reduced rate of *f* * *k* where ‘*f*’ denotes the proportion of translation occurring in the presence of CHX, factoring in the reversible nature of CHX inhibition. Newly translated GFPs are present in their non-fluorescent state, called immature GFP. The immature GFPs undergo post-translation folding and a 1-step maturation process via the oxidation and dehydration reactions between the conserved residues Y66, G67, R96, and E222 to form the fluorescent GFPs (mature) [64]. The level of immature GFP in a cell at a given point of time is denoted as *GFP*_*im*_(*t*), and the level of mature GFP in a cell is denoted as *GFP*_*m*_(*t*). Thus, after transcriptional and translational shut-off, at a given time *t*, the cells contain a mix of immature and mature GFP (**Figure 1B**). The equations **1** and **2** are the ordinary differential equations (ODE) quantifying the changes in mature and immature GFP proteins (*see materials and methods/Model*. The set of ODE is solved using the steady state GFP levels (**eq.3** and **eq.4**). The resulting solution (**eq. 5**) is the normalized GFP intensity. We fitted the single-cell GFP trace data to the normalized GFP intensity equation (**eq. 5**) using the Optimx() function in R [65, 66](*see materials and methods/Model fitting*) and estimated the parameters of the model. The *f* parameter measures the leakiness of translation due to cycloheximide’s reversible nature of cycloheximide [67] on translation elongation. A value of *f* = 0 indicates CHX is 100% effective in blocking translation, and *f* = 1 indicates no translation inhibition. We find that the median value of the *f* parameter across all replicates is 0 with a standard deviation of 0.021 for yeGFP-mODC and 0.04 with a standard deviation of 0.058 for the yeGFP-CLN2 degron as estimated by (**eq. 5, Figure 2C**). This suggests we had potent inhibition of new GFP protein synthesis under the experimental conditions used. The median rate of maturation (*µ*) is 0.58 *min*^*−*1^ for the yeGFP-mODC degron and 3.25 *min*^*−*1^ for the yeGFP-CLN2 degron (**Figure 2D**). For the cells with a very high rate of maturation (1 *min*^*−*1^), maturation is considered to be instantaneous and inconsequential to the decay kinetics of GFP. The median single-cell decay rate was 0.1 *min*^*−*1^ for yeGFP-mODC and yeGFP-CLN2 degrons (**Figure 2E**). The decay rate was used to estimate the single-cell half-life of degron GFP (**eq. 6**). The single-cell half-lives follow a gamma distribution with a positively skewed tail (**Figure 2F**).

### Quantifying noise in protein decay

After estimating the single-cell decay rates, we wanted to evaluate noise in the process of protein decay. We observed a 5-fold range in the half-lives between the slowest and the fastest decaying degron GFP at the individual cells. We calculated the single-cell heterogeneity in the half-lives of degron GFP as the coefficient of variation (*CV*) defined as standard deviation over the mean. We find very similar CV values for both the degron GFPs - 0.23 (n= 1413) for yeGFP-mODC and 0.23 (n = 1248) for yeGFP-CLN2, (**Figure 3B**). The *CVs* observed in our study are in the lower end of the range of coefficient of variation (0.2-0.4) in single-cell half-lives of mammalian proteins reported by Alber et al. [38].

**Figure 3.**
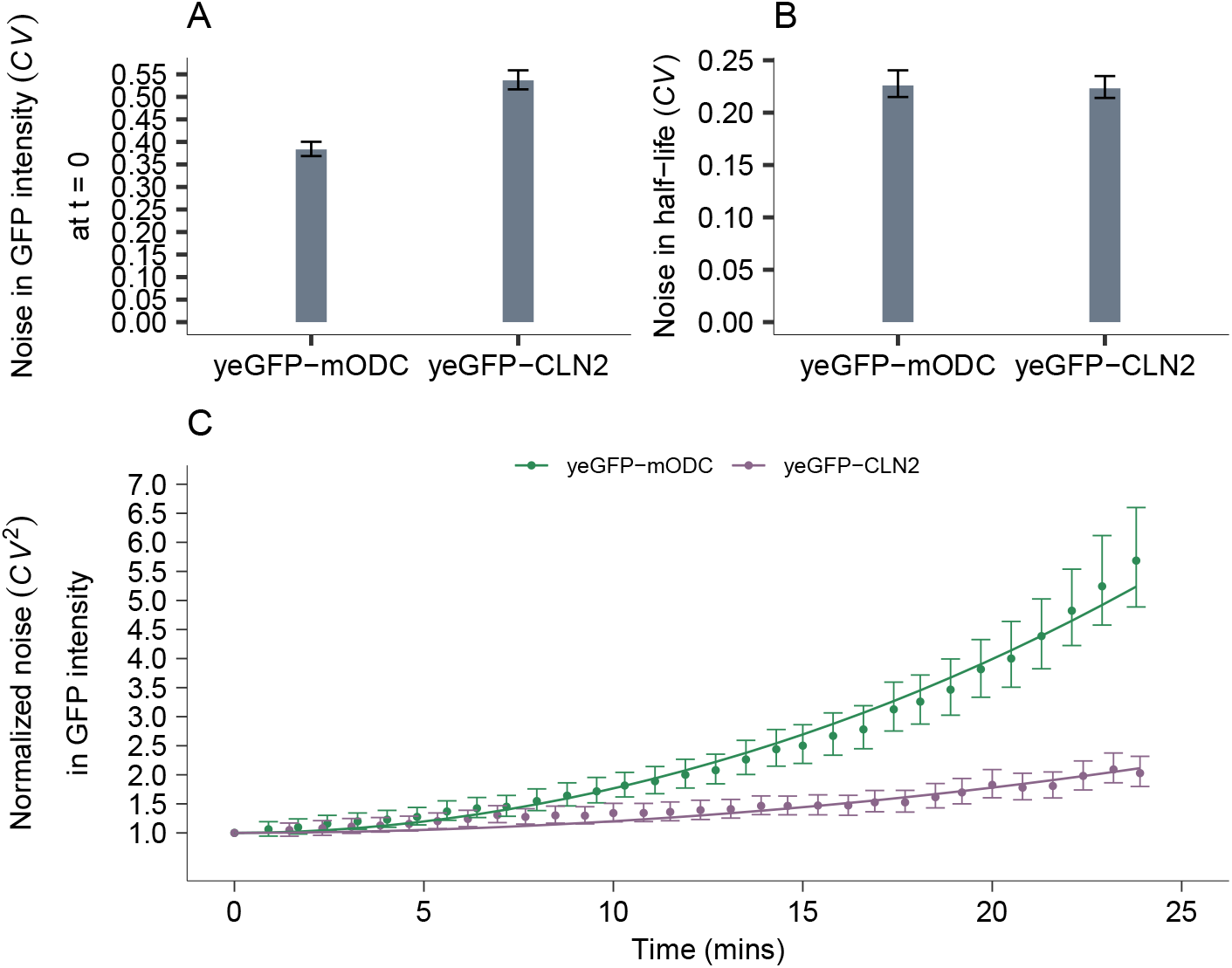
Cell-to-cell variability in the degron GFP half-lives. **A:** Expression noise in the degron GFP intensities. **B:** Noise in the decay rate of the two degron GFPs. Noise is calculated as 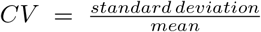. **C:** Changes in noise in GFP intensities over time. Noise is calculated as 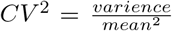 normalized to noise at *t* = 0. The solid lines represent the model fit of transient noise using **eq. 10** with the mean decay rate *m*_*d*_ for each degron GFP as 0.1 (shown in **Figure 2E**.), the *CV*_*d*_ in decay rates of 0.21 and 0.23 for yeGFP-mODC and yeGFP-CLN2, respectively. The fit resulted in an noise in GFP intensity at *t* = 0 (*CV* (0) = 0.25) for yeGFP-mODC and (*CV* (0) = 0.62) for yeGFP-CLN2. The error bars represent the 95% confidence interval from performing 2000 bootstraps on *CV* ^2^(*t*) normalized to the *CV* ^2^(0).

We wanted to confirm that the observed noise (*CV*) in the half-lives (thus decay rates) was biologically relevant and not due to measurement errors or parameter estimation. One way to assess whether the single-cell variability in the decay of the degron GFPs is due to biological heterogeneity is to evaluate the noise in GFP intensity over time after the perturbation of transcription and translation. Previous studies have shown that trends in noise in mRNA [37] and protein molecules [68] over time after perturbation of transcription reveals signatures of noise due to the processes active after perturbation. Since we block transcription and translation at the start of the time-lapse experiment, any changes in noise in GFP intensity during the time-lapse should be due to noise in protein decay. If the decay process is biologically noisy or stochastic, the noise in GFP intensity should increase monotonically over the time-lapse duration [69]. If there is no biological variability in the decay of the degron GFPs, i.e., *CV* in decay rates are closer to zero, noise (*CV* ^2^) in GFP intensity over time should remain constant throughout the time-lapse experiment. The initial (at *t* = 0) noise in GFP intensities for both the degron GFPs are shown in (**Figure 3A**). The change in GFP intensity noise relative to the expression noise at t = 0 is shown in (**Figure 3C**). The relative or normalized noise in GFP intensity increases monotonically for the degron GFPs. The increase is more prominent for yeGFP-mODC compared to yeGFP-CLN2.

Furthermore, a function for transient noise (*CV* ^2^(*t*), **eq. 10**, *see materials and methods/Estimating transient noise in GFP expression*) derived from a stochastic model of gene expression where the protein degradation fluctuated stochastically, and the protein decay rates were assumed to form a gamma distribution, explains the trend in noise in GFP intensity over time, proving that the change in noise in GFP intensity is due to variability in protein decay. This collectively proves that the observed noise in the single-cell decay rates (or half-lives) of degron GFPs is biologically relevant and not a result of an error in measurement or single-cell parameter optimization.

### Noise in protein decay is due to decay via the proteasomes

Proteolysis in yeast occurs via two different mechanisms. The proteasomal machinery, or the proteasome, degrades unstable or short-lived proteins, whereas most stable and misfolded proteins decay in the vacuoles [70]. Furthermore, cellular stress due to changes my pH, heat shock, and nutrient deficiency can cause proteins to degrade in vacuoles [71, 72]. We wanted to confirm that the noise observed in the decay of degron GFPs was due to decay via the proteasomes. If the proteasome decays the degron GFP, proteasome-specific inhibitor treatment should increase the overall median half-lives of the degron GFP. On the other hand, if degron GFPs are decaying through the vacuoles, the proteasome-specific inhibitor drug should be no change in the half-lives of the GFP. Thus, we treated cells expressing yeGFP-mODC degron with a proteasome-specific inhibitor MG132 for 30 mins before imaging. The proteasome inhibitor treatment resulted in a slower decay of the yeGFP-mODC degron, and this response was dose-dependent (**Figure 4 A and supplemental figureS3 A-B**). The 30-minute inhibitor treatment also resulted in a higher GFP intensity at the beginning of the time-lapse experiment than the control (0.1% DMSO) (**Figure 4B**). There was no change in the induction of GFP from the *GAL1* promoter between the control and drug treatment. Therefore, the increased GFP intensity at the end of the 30min treatment with the proteasome inhibitor compared to the 30min treatment with 0.1% DMSO control indicates an increase in the stability of the degron GFPs and not a higher expression of GFP.

**Figure 4.**
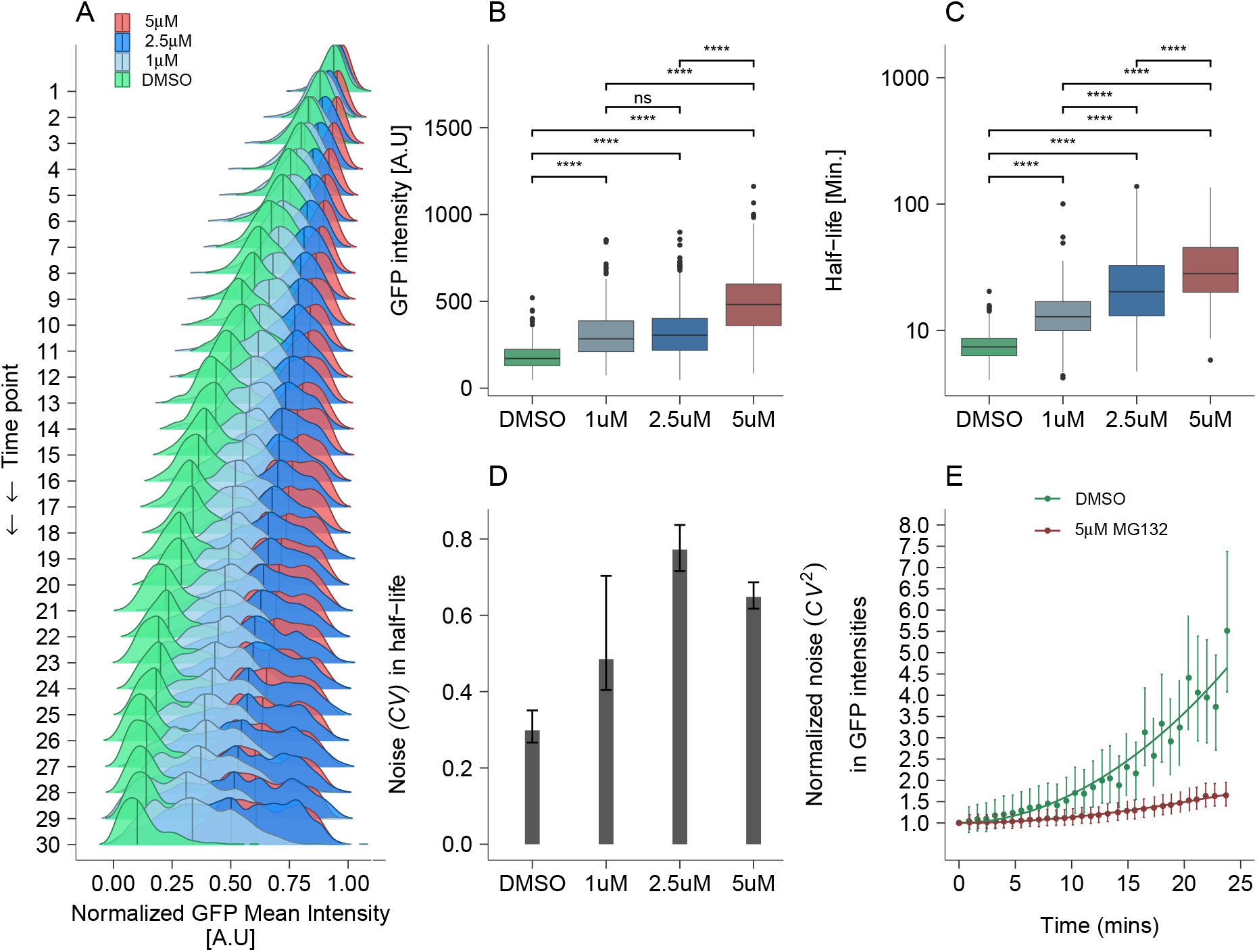
Effect of proteasome inhibition on cellular heterogeneity in the degradation of yeGFP-mODC. Cells were treated with the proteasome inhibitor MRG132 at varying concentrations for 30 mins after 2.5 hrs of degron GFP induction. 0.1% DMSO treatment was used as the control. **A**: Bulk representation of the reduction in GFP intensity over the time series. The x-axis corresponds to the distribution of GFP intensities at time *t* normalized with the GFP intensity of the cell at the first time point. Different colors correspond to the 0.1% DMSO control and MG132 drug treatment at 1*µ*M, 2.5*µ*M, and 5*µ*M concentrations. **B**: The mean GFP intensity of the cellular population increased upon the drug treatment. **C**: The half-life of the yeGFP-mODC degron increased from 7 mins (DMSO control) to 29 mins (5*µ*M MG132) with the drug treatment. The significance of the comparisons depicted by the bars on top of the boxplots was calculated by performing a t-test. **D**: Noise (*CV*) in the estimated half-lives of the degron yeGFP-mODC in the presence and absence of the proteasome inhibitor MG132. The error bars represent the 95% confidence interval calculated by performing 2000 bootstraps. **E**: Noise in GFP intensity over the 25 mins of the timelapse. Noise in GFP expression, calculated as 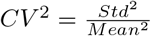, was normalized to noise at the first time point. The error bars represent the 95% CI of the 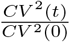 calculated by performing 2000 bootstraps. The noise increased monotonically in the DMSO control, whereas the noise over time became flat with increasing concentration of MG132 drug. The solid lines represent the model fit of transient noise using **eq. 10** with the mean decay rate *m*_*d*_ for DMSO control as 0.095 *min*^*−*1^, and *m*_*d*_ for 5*µ*M = 0.026 *min*^*−*1^. The noise in decay rates *CV*_*d*_ for DMSO control was 0.26 and 0.53 for the 5 *µ*M MG132 treatment.

Additionally, we observed an increased median half-life of yeGFP-mODC (**Figure 4C**) in the drug treatment group compared to the control. The increase in the median half-life of the degron is in line with the expected result if the proteasome degraded the protein. These results collectively show that the decay mechanism of yeGFP-mODC is via the proteasome and not by vacuoles.

We also wanted to evaluate the effect of proteasome inhibition on the noise in the decay process. First, the inhibition of the proteasome increased the cell-to-cell variability in the half-lives of yeGFP-mODC (**Figure 4D**). After perturbation of transcription and translation, fluctuations in the decay process result in a monotonic increase in GFP intensity noise relative to the initial expression noise in GFP intensity during the time-lapse (**Figure 3C, Figure 4D**). The proteasome inhibitor treatment lowered the extent of relative GFP intensity noise increase (**Figure 4D and supplemental figure S3C**). The lack of change in GFP intensity noise relative to the initial noise makes intuitive sense since if there are no fluctuations in the proteasomal decay during the time-lapse experiment, the GFP intensity noise should not change throughout the experiment. These results help solidify two main points. We can successfully study the proteasomal decay of yeGFP-mODC using our experimental setup, and the observed single-cell heterogeneity in the decay rates is due to noise in proteasomal decay.

### Cellular features explaining the cell-to-cell variability in GFP decay rates

Given the observed cell-to-cell variability in protein degradation of the two degron GFPs, we wanted to assess what cellular features contribute to this variation. Cell cycle stage and cell size are the predominant predictors of single-cell heterogeneity in mRNA expression [29]. Hence, we examined if the cell cycle stage can explain the variability in the decay rates of GFP. Synchronizing cells to a specific stage using hormones and drugs can mitigate the variability of cell-cycle stages. These treatments, while arresting cells to a particular cell-cycle stage, cause various physiological changes [73–75], making it difficult to distinguish the impact of the treatment itself from that of the cell-cycle stage on the single-cell variability in decay rates. Due to this, instead of using cell-cycle arrest to synchronize cells, we used the cell’s area, cellular shape, and DNA content as a proxy for the cell cycle stage. We define the cellular shape as the ratio of the cell’s short axis to the cell’s long axis. Cells at the beginning of the G1 phase are more circular. Thus, this ratio is closer to 1 for cells in the G1 phase [76]. To assess the DNA content of the cell, we used a Hoechst 33342 DNA stain. The dye preferentially stains DNA and is concentrated in the Nucleus of live cells. We used the cell’s area to calculate the cell size. Since the single-cell variability in the half-lives of the degron GFPs was due to noise in proteasomal machinery, we also assessed if the number of proteasomes in a cell dictates the cell-to-cell variability in the degron GFP half-lives. For this, we tagged the catalytic *β*-subunit of the 20s core protein of the proteasome named Pup1 with a tDimer red fluorescent protein and quantified the mean intensity per cell. Tagged Pup1 protein was localized in a puncta in the cell and colocalized with DNA stain consistent with previous findings [77] (**Supplemental figure S4**). We quantified the intensity of pup1-tDImer in a cell as the average intensity of the puncta in the cell. Lastly, we also evaluated if the amount of GFP in a cell at the initial time point played a role in the decay dynamics of the GFP in that cell.

To evaluate the relationship between each cellular feature and the estimated single-cell half-lives of the degron GFPs, we first looked at the individual scatterplots and simple linear regression between the single-cell half-lives of the degron GFPs and the cellular features (**Figure 5A**). The cellular shape did not correlate with the degron GFP half-life in that cell. The area of the cell and the DNA content correlated positively with the single-cell degron half-life, irrespective of the degron GFP. The relationship was stronger for yeGFP-CLN2 degron as compared to the yeGFP-mODC. The amount of GFP intensity at *t* = 0 of the time-lapse experiment had a significantly positive correlation with the degron GFP half-life. This is intuitive as cells with a higher half-life (lower decay rate) of the degron GFP will have more GFP molecules in the cell. Based on the 2D scatter plots, the amount of Pup1 in a cell did not correlate with the degron GFP half-life of the yeGFP-mODC. On the other hand, the amount of Pup1 showed a small, albeit significant, positive correlation with the yeGFP-CLN2 half-lives. Both of these results were surprising as the expectation is that the amount of proteasomal machinery in a cell should negatively impact the half-life of the degron GFPs. To further investigate this, we examined the relationship of the amount of Pup1 with other cellular features. The Pup1 intensity in a cell scaled with the area of the cell (**Supplemental figure S5A-B**) and with the amount of DNA in the cell. The amount of DNA in the cell also correlated with the cell size (**Supplemental figure S5C**). Since these cellular features are highly correlated, it becomes harder to decipher the true relationship of these features with the single-cell half-life of the degron GFP. To remove the effect of the correlation of area, DNA, and Pup1 with each other on the correlation of each with the degron half-lives, we calculated the partial Pearson correlation of the cellular features (area, amount of Pup1, DNA, and GFP) of a cell with the half-life of GFP in the cell. The partial Pearson correlations compared to standard pairwise Pearson correlations from the scatterplots are shown in (**Figure 5B**). When controlled for the area, amount of DNA, and GFP in the cell, the amount of Pup1 in a cell has a significant negative (albeit small) correlation with the degron GFP half-lives. This correlation is even smaller for yeGFP-CLN2-expressing cells. The partial correlation between the GFP intensity of the cell and the half-life of GFP in that cell when controlled for other cellular features did not change as compared to the standard Pearson correlation. This is because the GFP intensity of a cell did not scale with cellular features of the cell (**Supplemental figure S5D**). Cell’s area showed a significant positive partial correlation with the half-lives of the degron GFP. The relationship is larger for the yeGFP-CLN2 degron as compared to the yeGFP-mODC. Bigger cells had more total DNA stains of the nucleus (**Supplemental figure S5B**), representing cells further along in the cell cycle progression [78, 79]. If the area of the cell is used as a proxy for the cell cycle stage, considering how cells in the G1 stage are smaller and have less DNA content compared to cells in later stages of the cell cycle, this result makes sense. G1 cyclin, Cln2, is unstable in G1 cell cycle stage [80]. Hence by using the area as a proxy for the cell cycle stage, smaller cells, more likely to be in the G1 phase, have a lower half-life of the yeGFP-CLN2 degron.

**Figure 5.**
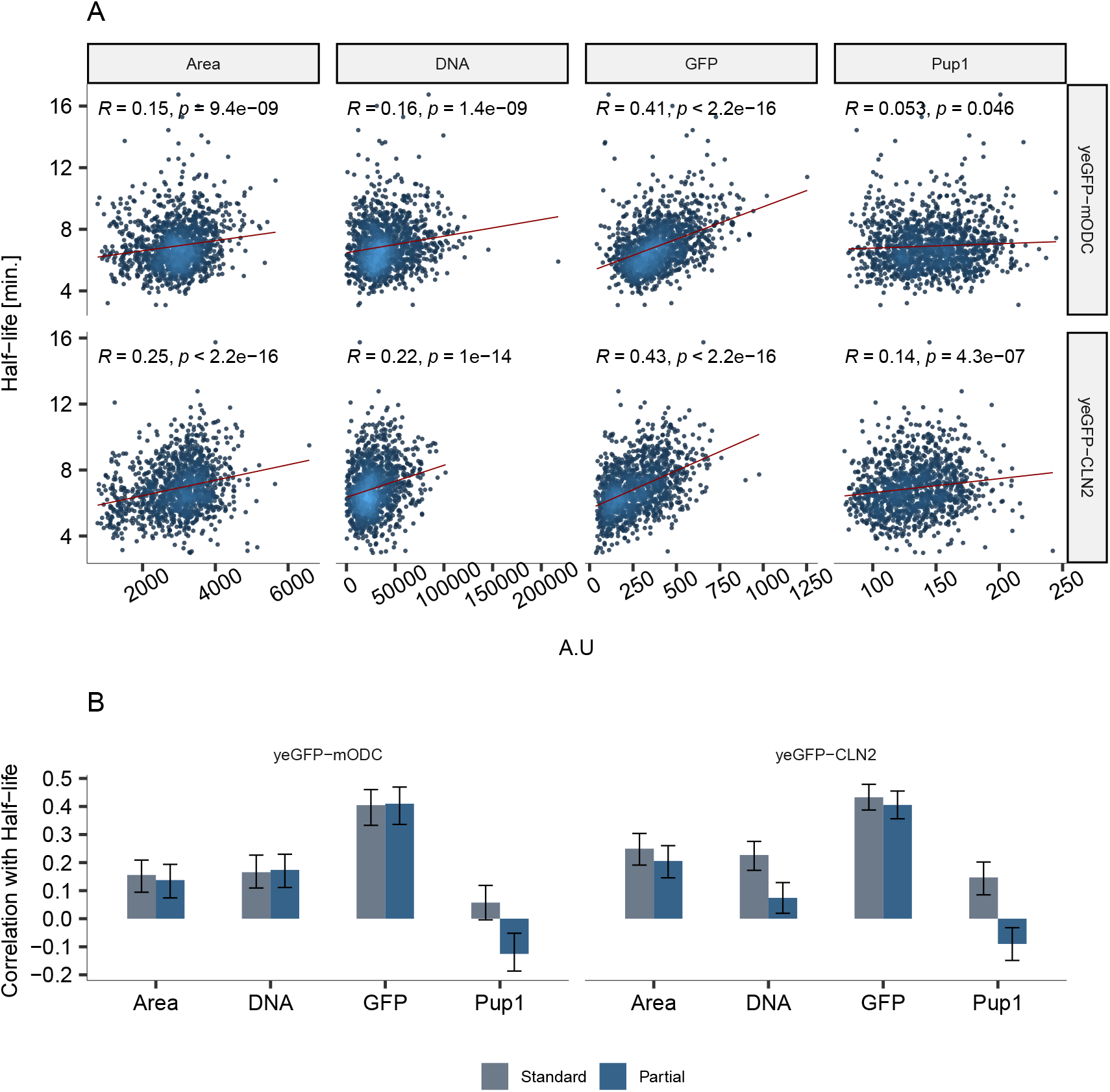
Cellular features explaining the cell-to-cell variability in the degron GFP half-lives. **A**: Pearson correlation between cellular features and the half-lives. Each point corresponds to a single cell. The area of the cell is calculated as the sum of the triangles enclosed within the surface mask of the cell. DNA refers to the total Hoechst 33342 intensity of the cell’s nucleus. GFP column is the mean GFP intensity of the cell. Pup1 is the mean intensity of the tDimer-RFP puncta within the cell. All the values of the cellular features are from the first snapshot from the time-lapse series. **B**: Comparison of the standard Pearson correlation and the partial Pearson correlation of the GFP half-lives with the cellular features. Area, the DNA content of the cell, and the amount of Pup1 in a cell are correlated (**Supplemental figure S5A-C**). A pairwise partial correlation was performed between each cellular feature and the degron GFP half-life of the cell using the pcor() function. The gray bars correspond to the standard Pearson correlations, and the blue bar corresponds to the partial Pearson correlations calculated from cor() and pcor() functions in R, respectively. The error bars represent the 95% confidence interval from performing 2000 bootstraps.

## Discussion

Noise in gene expression can arise from each of the biochemical steps involved in the synthesis and decay of mRNA and protein molecules in the cell. Until recently, studies evaluating gene expression noise have focused primarily on sources involved in synthesizing mRNAs and proteins by estimating the transcription kinetics from their abundances at a single time point. While these studies have led to some significant understanding in the field, studying the variability in the kinetics of the underlying processes of gene expression can help us better understand the sources and the modulation of noise.

In this study, we estimated the single-cell kinetics of the proteasomal decay of degron GFPs in thousands of cells following two different pathways to the proteasome. yeGFP-mODC degron GFP gets targetted to the proteasome independently of the ubiquitin ligation system, while yeGFP-CLN2 degron GFP gets targetted to the proteasome via the ubiquitin ligation pathway. The cell-to-cell variability in the single-cell decay rates of the two degron GFPs observed in our study, along with single-cell variability in the decay rates of mammalian proteins [38], shows that the protein decay process is substantially variable amongst an isogenic population of eukaryotic cells. Using the noise decomposition framework [30],(*See Materials and methods/GFP expression noise decomposition*), we find that the single-cell half-lives of the degron GFPs explain approximately 16% and 20% of the steady state variability (*CV* ^2^) in the expression of the yeGFP-mODC and yeGFP-CLN2 degrons. We also showed a considerable contribution of noise in protein decay to transient noise in GFP expression over time, signifying the variability in the process of protein decay. While the yeGFP-mODC and yeGFP-CLN2 exhibited similar noise (*CV*) in the single-cell decay rates (**Figure 3B**), protein degradation of each degron exhibited different degrees of influence on noise in GFP intensity after the perturbation of transcription and translation. We confirmed that yeGFP-mODC decays via the proteasome by treatment with a proteasomal inhibitor MG132. Since the two degron gets targetted to the proteasome via two different pathways, the difference in the contribution of protein decay on noise in GFP intensity over time shows different noise levels in each decay pathway. The yeGFP-mODC decay by the proteasome is ubiquitin independent. The higher slope of the noise in GFP intensity over time for yeGFP-mODC than yeGFP-CLN2 indicates a possible higher noise level in the ubiquitin-independent proteasomal decay. On the other hand, the low noise slope in GFP intensity over time for yeGFP-CLN2 indicates a possible lower noise level in ubiquitin-dependent proteasomal decay.

We also evaluated the contribution of cellular features to the variability in the instability of the degron GFPs. The amount of catalytic subunit of the proteasome explains only a small fraction of the observed variability in half-lives of the degron GFPs. This was surprising as the expectation was that the number of proteasomes in a cell, in general, should explain the variability in the decay of proteins. On the one hand, a possible explanation behind our observation is that proteins associated with the proteasome are less noisy [81], hence might not play a huge role in influencing the variability in the decay of proteins. However, the Newman et al. study did not include the Pup1 subunit measured in our study. On the other hand, the extremely weak correlation between the Pup1 subunit of the proteasome and the half-lives of yeGFP-CLN2 might indicate that the catalytic subunit might not be the rate-limiting factor in the decay of the degron. Since the PEST sequence in yeGFP-CLN2 degron belongs to the Cln2 protein (*see Materials and Methods*), the degron’s decay is controlled by similar mechanisms that regulate the rapid decay of the Cln2 protein [60]. The Cln2 G1 cyclin is self-limiting. The cyclin forms a complex with cdc28, a kinase promoting the initiation of the S phase. The CLn2-cdc28 complex activates the cdc28 kinase to phosphorylate the downstream substrates. The activated cdc28, in turn, also phosphorylates six amino acid residues in the PEST sequence on Cln2, marking it for rapid degradation [82] via the ubiquitination-dependent proteasomal decay. The phosphorylated C-terminal sites of the Cln2 protein are recognized by the E3 ligase, Grr1, of the SCF (Skp1/Cdc53/F-box) protein complex, that polyubiquitinates the protein [82–84] The E3 ligase, Grr1, is necessary for decaying the Cln2 protein [85]. Interestingly, the Grr1 protein is also degraded rapidly by the ubiquitin-dependent proteasomal decay, with a half-life of 20mins when grown in rich media [86]. Hence, the amount of Grr1 protein in a cell might dictate the variability in the half-life of the yeGFP-CLN2 degron. Similarly, we observed a minor but significant negative correlation between the amount of Pup1 in a cell and the half lives of yeGFP-mODC degron. This means other factors might have a more direct role in explaining the cellular heterogeneity in the decay rates of yeGFP-mODC degron. The regulation of decay of Ornithine decarboxylase (ODC), from which the degron sequence in yeGFP-mODC originates, happens via negative feedback. The decay of ODC depends on the cellular concentration of polyamines, like spermidine and spermine [87, 88]. The ODC stimulates the biosynthesis of polyamines. A higher concentration of polyamines induces the transcription of antizymes (Az1), which stimulates the decay of ODC by increasing the recognition of ODC by the proteasome. Although higher concentrations of antizymes increase the ODC decay, the decay rate remains unaltered by increased Az1 concentration [89]. The variability in the concentration of polyamines in a cell might explain the cellular heterogeneity in the yeGFP-mODC degron half-lives. Additionally, growth rates and cell density of culture also dictate the concentration of polyamines [90, 91], adding another layer of complexity to the regulation and, thus, noise in the decay of ODCs.

It should be noted that we considered a simple model of protein decay where both the mature and immature fluorescent protein decays at the same rate. While this is an appropriate approximation for fluorescent proteins, this is only sometimes true for endogenous proteins that form complex secondary structures. For instance, the endogenous yeast ODC exists as a dimer, and the immature (monomer) and mature (dimer) ODC have different stabilities [87]. This causes a non-exponential protein decay, where newly made proteins decay faster than the old proteins [49]. A considerable proportion of the eukaryotic proteome exhibits non-exponential decay [49]. On a similar note, a recent study has highlighted the necessity for modeling complex multi-step degradation of mRNAs to explain the observed sub-Poissonian noise in constitutively expressed proteins in fission yeast [92]. Hence, given the cell-to-cell variability observed in proteasomal decay in our study and the time-dependent decay of some endogenous proteins [49, 50], it is essential to model gene expression with complex decay dynamics (as opposed to deterministic exponential decay) to truly estimate the role of each process in studying noise.

Furthermore, our study highlights the importance of estimating the single-cell kinetic parameters of gene expression instead of estimating a single kinetic rate for gene expression processes. Inferring kinetic parameters for an isogenic population of cells from steady-state distribution of mRNA and protein molecules ignores the differences in rates as a noise source. As argued in this study and elsewhere [52, 93], single-cell kinetic rates vary in a population of isogenic cells, and one should not ignore this cellular variability in rates while studying gene expression noise.

## Materials and Methods

### Construction of the GFP expressing plasmids

The two degrons in this study are previously well characterised degron GFPs. The yeGFP-mODC degron contains the mODC PEST sequence that is 28 amino acids long, which targets the protein to the proteasome in a ubiquitin-independent manner [58]. The yeGFP-CLN2 degron contains the last 180 amino acids from the C-terminal of the Cyclin 2 protein. This sequence comprises 37 amino acids, containing the PEST sequences and other residues necessary for the rapid degradation of the Cyclin 2 protein via the ubiquitin pathway [62]. Briefly, the degron GFP expression cassette was designed to be under the transcriptional control of the galactose inducible promoter, *Gal1*, and the expression cassette was genomically integrated into the *LEU2* locus of the yeast strains. A detailed description of the construction of the GFP-expressing cassettes is provided in the Supplemental methods

### Yeast strains

All the strains in the study were made on the MRG 6301 background (*Mata ADE2 can1-100 his3-11,15 leu2-3,112 trp1-1 ura3-1*), which Dr. Marc Gartenberg provided. The strain SDY10 was created where the *BAR1* gene was deleted from the parental background via homologous recombination using PCR-amplified SpHis3 region from the pFA6a-HIS3MX6 plasmid with SD26 and SD27 primers. For the Pup1 quantification, the SDY11 strain was created where the endogenous Pup1 gene was tagged with tDimer-RFP using an integrating plasmid LEP771 provided by Dr. Kiran Madura. Briefly, the plasmid was linearized with the EcoNI restriction enzyme and transformed into the SDY10 strain. The plasmids expressing the degron GFPs (yeGFP-mODC and yeGFP-CLN2) were genomically integrated at the *LEU2* locus of the SDY11 strain via homologous recombination with the BstEII linearized constructs. Transformants were screened for single and multiple insertion events using PCR primer pairs. Only strains with a single integration event were chosen for the study. The strain details are provided in the 3, and all the PCR primers used in the study are listed in 1.

### Yeast growth and media

Cells were streaked out on YPD plates. Single well-separated colonies from YPD plates were grown for 4 hrs in YPD liquid media. Overnight cultures were prepared by diluting the YPD cultures in 2ml synthetic complete (SC) media without Uracil and Leucine (SC-ura-leu) in 1:300 dilution. The SC-ura-leu media was supplemented with 2% raffinose and 0.1% glucose as the carbon source for overnight growth. The 1:300 dilution in SC media results in exponentially growing cells in 12-14 hrs (from discussions with Dr. Marc Gartenberg and Dr. Melinda Borrie). Cells growing exponentially in raffinose media result in rapid induction with galactose [94] and reach a steady state of expression faster than cells growing in glucose media. The cells were then diluted to OD 0.1-0.2 the next day in SC-ura-leu media supplemented with 2% galactose. Cells were grown for 3 hrs in the galactose media to induce GFP. All the cultures were grown at 30C with shaking at 200 rpm.

### Proteasomal inhibition

Proteasomal inhibition via drugs like MG132 requires the usage of mutant strains like *erg6*Δ and *pdr5*Δ [70] to increase the cellular permeability, retention, and the cellular concentration of the drug. These mutations can lead to physiological changes in the cells making the direct interpretation of the proteasome inhibition on cellular heterogeneity in decay rates harder. To avoid this, we adopted a non-genetic approach to increase the susceptibility of *Saccharomyces cerevisiae* to MG132 [95]. Briefly, the cells from single colonies were grown in YPD media, then diluted in SC-ura-leu + 2% raffinose + 0.1% glucose media, where proline was used as the nitrogen source instead of ammonium sulfate.

This was achieved by using a yeast nitrogen base (YNB) without ammonium sulfate and adding 0.1% proline to the media. After an overnight growth in this media, cells were diluted into SC-ura-leu + 0.1% proline + 2% galactose media supplemented with 0.03% SDS to facilitate the transient opening of the cell wall. After 2.5hrs of growth in the above media, cells were either treated with empty vehicle (0.1% DMSO) or 1*µM*, 2.5*µM* and 5*µM* of MG132 dissolved in 100% DMSO. Cells were grown for 30 mins and then imaged in the presence of the treatment (DMSO or MG132).

### Microscopy

#### Preparing cells for microscopy

After 3 hrs of growth in SC-ura-leu media with galactose, cells were sonicated (1 sec on 1 sec off pulse for 20sec with 40-60% amplitude), spun down (quick spin at 8000g), and concentrated appropriately before plating on 96-well glass-bottom plates (from Ibidi, cat. No. : 89621) which were coated with concanavalin A (cat. No. J61221). This prevented cells from moving while imaging. 200*µl* of the concentrated cell culture were plated onto individual wells for 10 mins, the wells were washed with sterile water twice and replaced with fresh SC-ura-leu + 2% galactose media till each field of view was selected for imaging. Two drops of NucBlue (cat. # R37605) was added to each well to stain the DNA of the cells. After selecting all fields of view, the SC-galactose media was replaced with SC media with 2% glucose and 100 *µg/ml* cycloheximide. The cells were imaged immediately afterward. The parental strain (SDY10) lacking any fluorescent tags was used as an autofluorescence control and was included in all the experiments.

#### Image acquisition

Each field of view was imaged for a duration of 20-25mins with each image being captured at 40-50 sec intervals. The time-lapse experiment was conducted at 30C. Microscope: Images were acquired using a Nikon TiE fluorescent microscope. The 96-well glass-bottomed plate was mounted onto the MA60 microplate holder attached to the TI-SAM attachable mechanical stage. Camera: The images were acquired using the Prime 95B sCMOS camera from Teledyne Photometrics using 1x1 binning. PFS: Cells were kept in focus during the time-lapse duration by using Nikon Ti-E’s perfect focus system (PFS). Differential Interference Contrast (DIC) images were acquired using the 6V 30W dia Pillar illuminator with an exposure time of 4ms. Onstage incubator: An external temperature-controlled incubator attachment was used to maintain the cells at 30C. Fluorophore channels: GFP fluorescent images were imaged with an exposure time of 40 ms, with 20% intensity, using 475 nm excitation and a 540 nm emission filter. Tdimer-RFP fluorescent images were imaged with an exposure time of 10 ms, with 20 % Xcite intensity, 545 nm excitation /640 nm emission. DAPI images were imaged with an exposure time of 50 ms, with 30% light intensity, 395 nm excitation, and 450 nm emission. All the image acquisition during the time-lapse for various fields of view was automated using the Metamorph software version 7.10.3.279. The raw image files are published on Dryad https://doi.org/10.5061/dryad.bnzs7h4g6

### Data analysis

#### Image processing

DIC images were used to segment single cells using the YeastSpotter tool [96]. The tool was implemented as a Python script using individual DIC images as the input. The cell segmentation of each image was parallelized using Rutgers’s advanced computing system. Each segmented cell was then tracked in all the images for a given time-lapse experiment using the surface identification and tracking features of the Imaris suite (version 9.8). Cells were assigned a unique identification number, and various cellular features like fluorescent intensities, area, sphericity, etc, were quantified. The pup1-tDimer intensity of the cell was calculated as the average intensity of the Pup1-tDimer punta in the cell. The nucleus was segmented using the Pup1-tDimer stain, and only this segmented nuclear region was used to quantify the total amount of Hoescht 33342 intensity of the cell. The background intensity of each image was calculated using an in-house ImageJ macros script. The exact time intervals between each time-lapse image were calculated using the image acquisition time extracted from the image’s metadata using a custom ImageJ macros script and R. All the downstream data processing of the single-cell intensities is done in R. The scripts for processing the images are submitted to the GitHub repository code/imageJ: https://github.com/shahlab/ProteinDecayNoise-paper.git

#### Single-cell fluorescent intensity calculations

The single-cell measurements from Imaris were read in R. Each cell is associated with a unique track ID, area, GFP mean intensity, tDimer-Red mean intensity for experiments with Pup1-tDimer, DAPI mean intensity, the time elapsed since the first image was taken, and the number of voxels. The mean fluorescent intensity of each cell is subtracted by the mean background intensity for that image. Dead cells were identified and removed from the analysis based on the live dead staining using NucBlue. Autofluorescence intensity was calculated based on the intensity of the parental strain SDY10 lacking fluorescent tags. The autofluorescence threshold was defined for each time point as the 95th percentile of the intensity distribution of the autofluorescence control. The autofluorescence intensity was subtracted from each cell at every time point. Cells with an intensity less than the autofluorescence intensity in the first time frame were excluded from the analysis. For subsequent time frames, single-cell intensities less than the autofluorescence intensity were replaced with NAs. Only cells with non-NA values for all 31 timeframes were included in the analysis. Furthermore, Cells with GFP intensity values higher than GFP intensity at the first time point in more than 5 images were excluded from the analysis since this might be due to technical issues, like cells going in and out of focus during the time-lapse or errors in tracking. Code under code/generate df dir in the github repository : https://github.com/shahlab/ProteinDecayNoise-paper.git

### Model fitting

#### Model

A mechanistic model of GFP decay developed is explained below. The galactose carbon source in the media induces the transcription of the GFP mRNA molecules, which gets translated into immature GFP molecules at the rate of *k*. The levels of immature GFP in a cell at a given time *t* are given by *GFP*_*im*_(*t*). Each immature GFP undergoes a one-step maturation process at the rate of *µ* to make the mature form of GFP. The amount of mature GFP in a cell at any given moment is denoted as *GFP*_*m*_(*t*). Right before the time-lapse imaging begins, the transcription of new GFP mRNAs is inhibited by replacing galactose with glucose as the carbon source, and the translation of new GFP molecules is inhibited by the addition of cycloheximide (CHX). Due to the reversible nature of translation inhibition by CHX, new immature GFPs are made at a rate of *k*\**f* where *f* is the degree of leaky translation due to the reversibility of CHX treatment and *k* is the rate of translation **1B**. These immature GFP then undergoes maturation at a rate *µ* to add to the pool of mature GFP *GFP*_*m*_(*t*). Both the mature and immature GFP can undergo proteasomal decay at rates *δ*_*im*_ and *δ*_*m*_, respectively. Hence, the rate of change of immature and mature GFP levels in a cell can be written as:

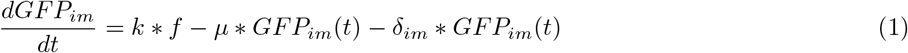

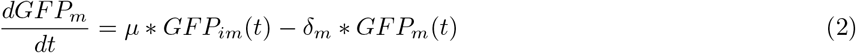

with the steady-state values of two GFPs as:

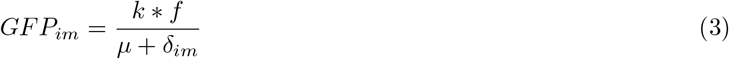

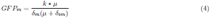

Assuming the mature and immature GFP decay at the same rate ie *δ*_*im*_ = *δ*_*m*_, now referred to as *δ*, the set of ordinary differential equations can be solved as:

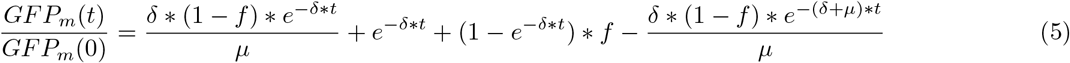

#### Parameter Estimation

The left-hand side represents the single-cell GFP intensity at time t normalized by the cell’s GFP intensity at t = 0. The single-cell GFP intensities were normalized with the GFP intensity from the first timeframe. The normalized GFP intensities were fitted to the **5**. The parameters were estimated for single cells by minimizing the residual sum of squared errors. The minimization was performed using the Optimx package in R, using the “L-BFGS-B” method [65, 66]. The bounding constraints on the parameters were:

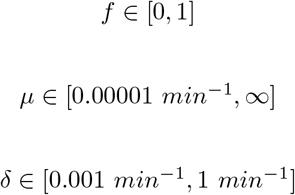

The half-life of a GFP was calculated as

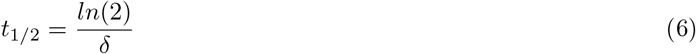

The bounding constraints on the parameter *f* for experiments with proteasome inhibitor MG132 (both treatment and control) were set to *f* ∈ [0, 0.1] since experiments without proteasome inhibitor in yeGFP-mODC expressing cells resulted in a median *f* value of 0.

### Estimating transient noise in GFP expression

Several works have tried to derive the transient noise after perturbing gene expression parameters [68, 97]. Following their derivation, we investigate the noise in GFP levels after the translation block, assuming that the first-order decay rate *δ* is drawn from a gamma distribution with mean *m*_*d*_ and coefficient of variation *CV*_*d*_. For this derivation, we assume that the initial level of GFP is independent of the decay rate.

The expected value of the mean GFP level is given by the moment-generating function of the gamma distribution

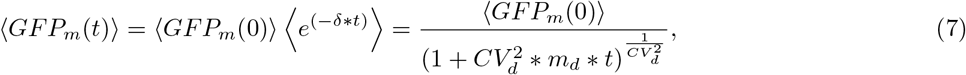

where ⟨*GFP*_*m*_(0)⟩ is the mean GFP intensity of the population at *t* = 0 and *CV* (0) is the coefficient of variation in *GFP*_*m*_ levels at *t* = 0. Similarly, the second-order moment is obtained as

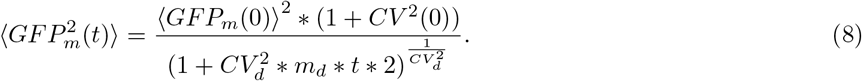

This leads to the following transient coefficient of variation in GFP levels

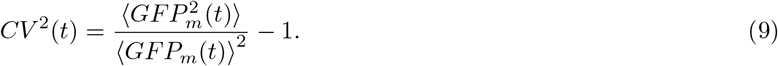

Substituting eq 7 and eq 8 in eq 9 results in the following *CV* ^2^(*t*) (normalized by the its value at time *t* = 0)

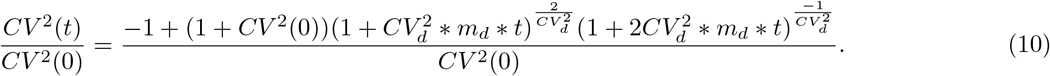

As shown in Fig. 3, depending on the extent of stochastic variability *CV*_*d*_ in the decay rate, *CV* ^2^(*t*) increases over time. In the limit *CV*_*d*_ → 0, this transient noise *CV* ^2^(*t*) → *CV* ^2^(0), and becomes invariant of time.

### GFP expression noise decomposition

Previous studies have formulated a framework to decompose the variance in the expression of mRNAs and protein expression into various sources like transcription, translation, and cellular volume [30]. Using the same framework, we estimated the noise in the steady state GFP expression due to variability in the half-lives of the degron GFP. We estimated the linear relationship of the GFP expression and the half-lives of the GFP in the same cell :

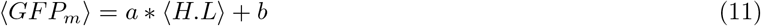

Where *GFP*_*m*_ is the mean GFP intensity of the cell at *t* = 0 and *H*.*L* is the *t*_1*/*2_ calculated from **eq.6**, *a* is the slope of the linear regression line and the *b* is the intercept.

Using these values, we estimated the squared *CV* in GFP intensities due to the GFP half-lives

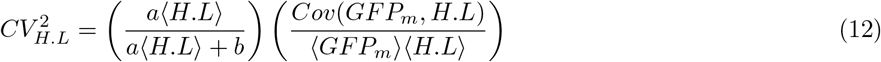

and calculated the percentage of noise in GFP intensity at the steady state due to the half-life of GFP as:

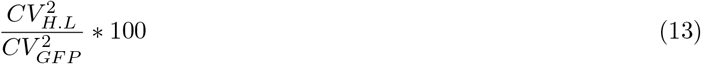

Where,

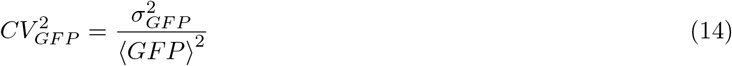

with *σ*_*GF P*_ is the standard deviation of GFP intensity and ⟨*GFP* ⟩ is the mean GFP intensity at *t* = 0.

## Supporting Information

### Supplemental Figures

**Figure S1.**
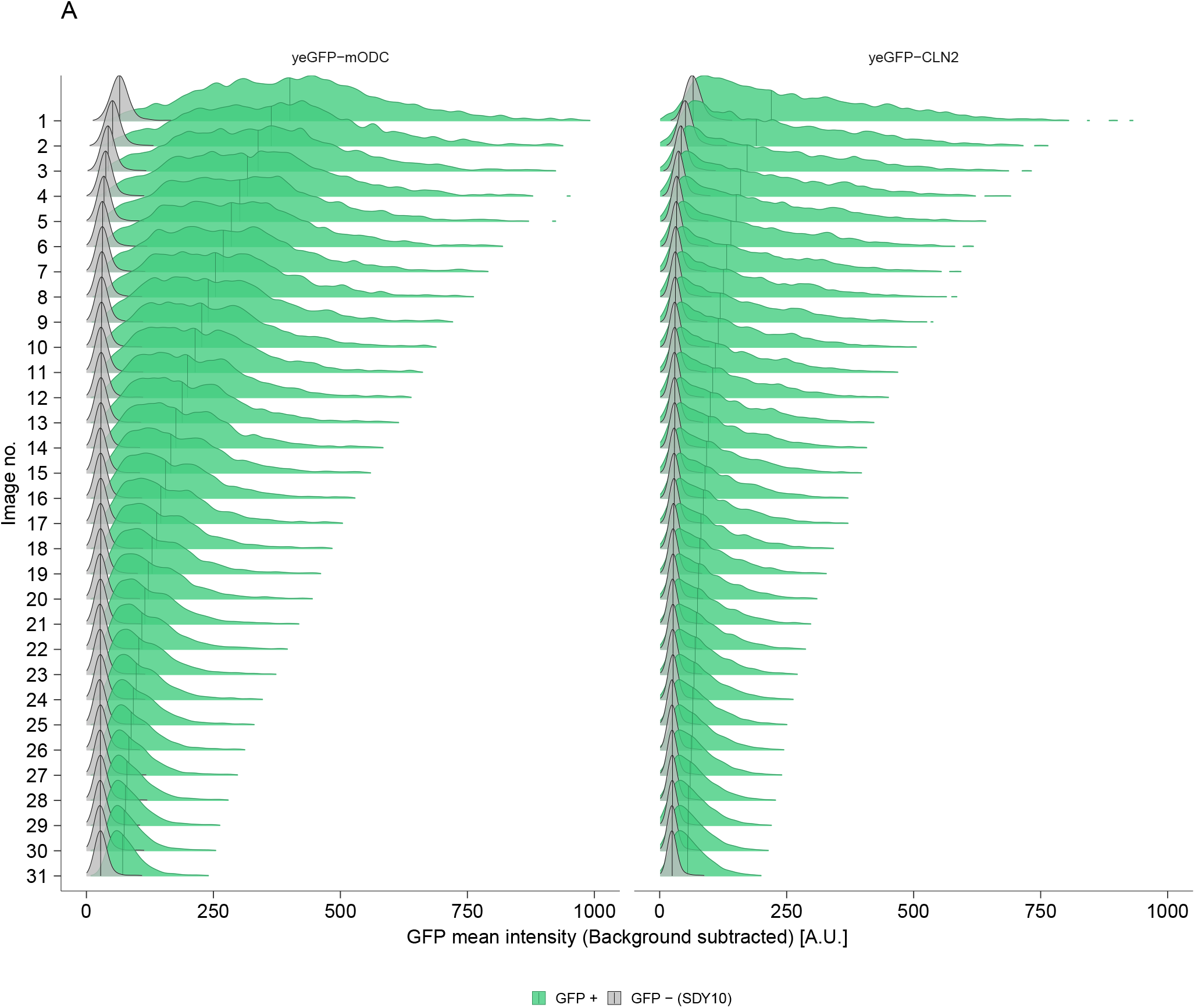
Autofluorescence and Positive GFP intensities. Raw changes in the mean GFP intensity of cell over the timelapse series. The gray distribution represents the autofluorescence intensities of the cells as calculated from the parental background strain, SDY10.

**Figure S2.**
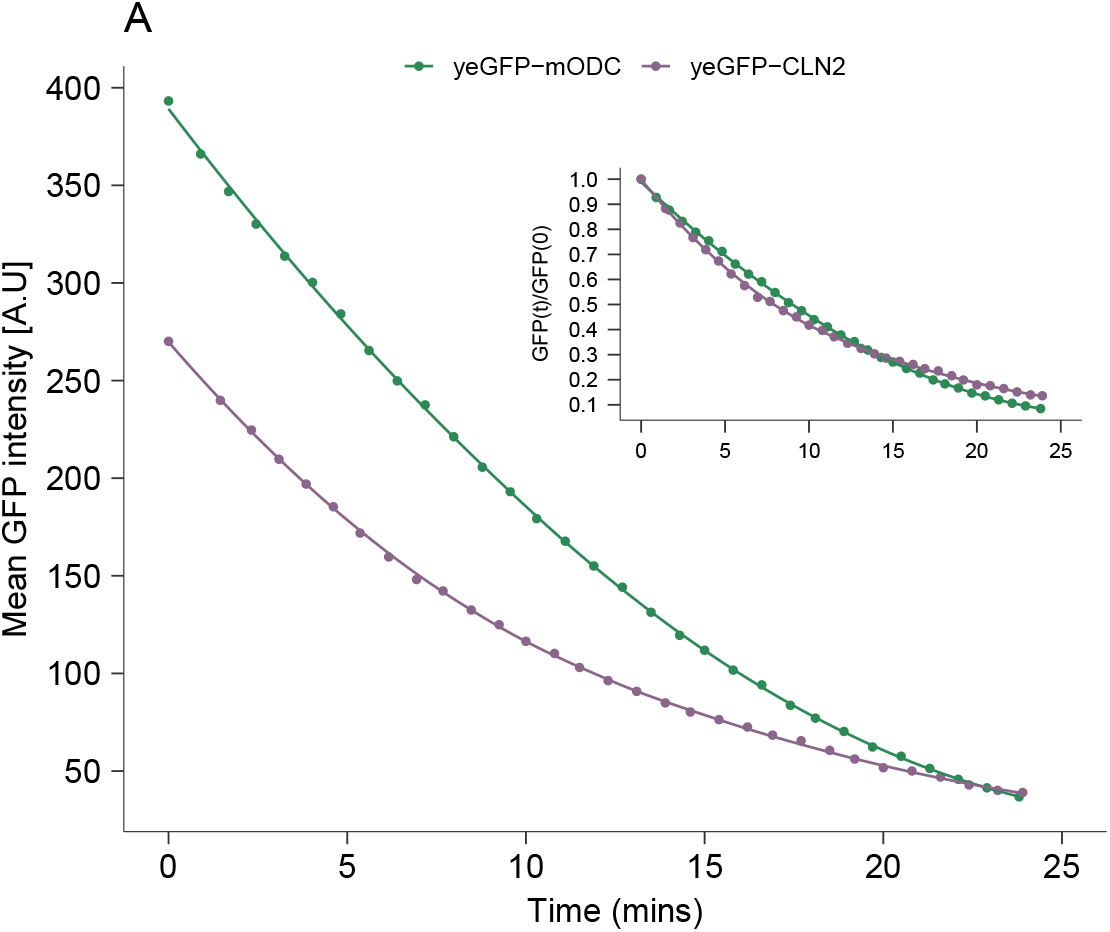
Reduction Mean GFP intensity. The mean GFP intensity of the population of cells is plotted against time. The two colors represent each degron GFP. The inset shows the relative change in mean GFP intensity over time.

**Figure S3.**
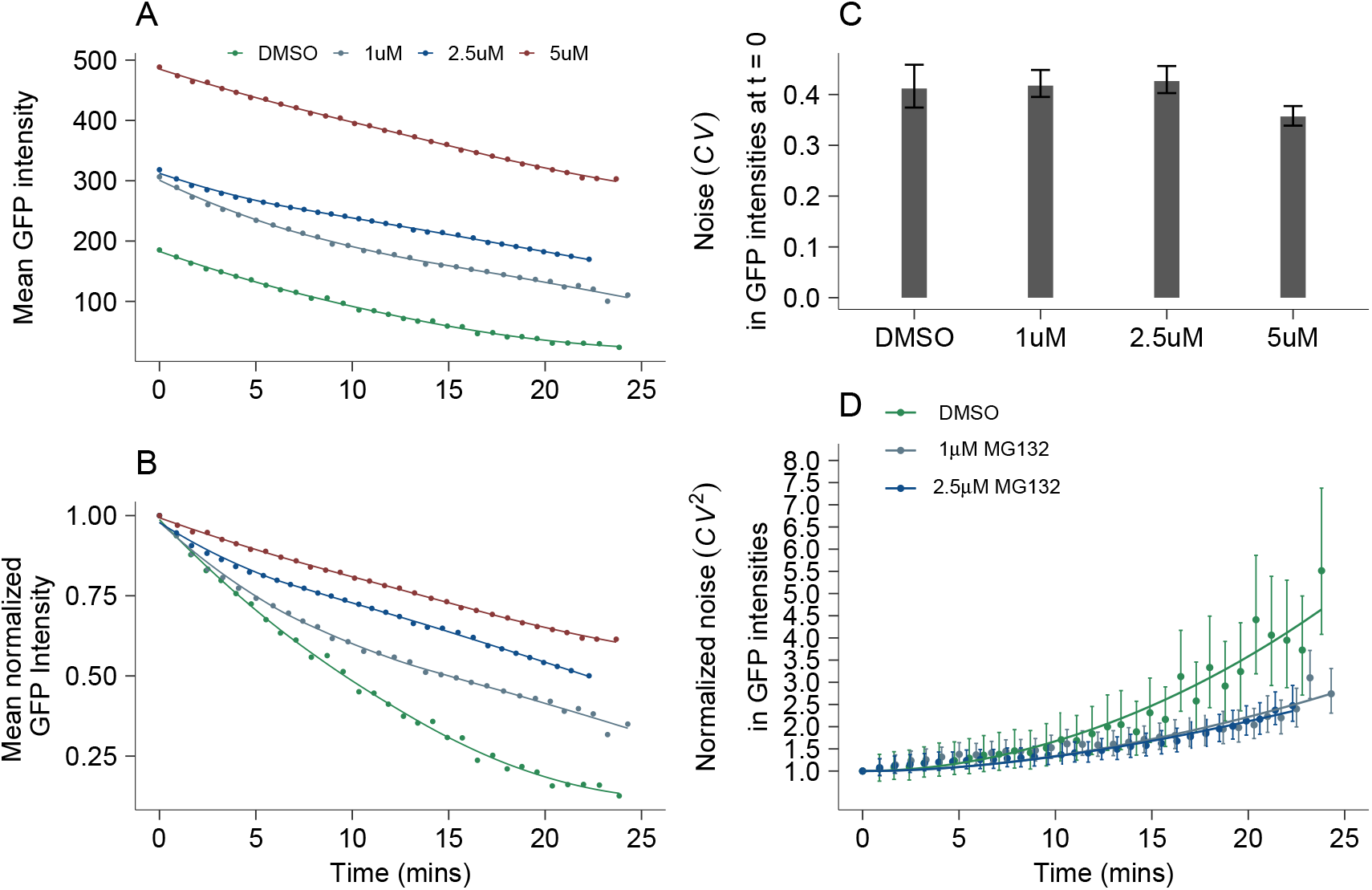
Effect of proteasomal inhibitor treatment on the decay of yeGFP-mODC. **A-B**: Mean GFP intensity trace (A) and relative change in mean GFP intensity (B) of yGFP-mODC treated with either control (0.1% DMSO) or with varying concentrations (1*µM*, 2.5*µM*, 5*µM*) of the proteasome inhibitor drug, MG132. **C**: Noise in GFP intensity at t = 0. The error bars represent 95% CI of *CV* by performing 2000 bootstraps. **D**: Relative change in noise in GFP intensity over time. Normalized noise 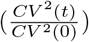 in GFP intensity is plotted at each timepoint after the perturbation of transcription and translation. The error bars represent 95% CI of 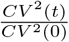estimated by performed 2000 bootstraps. The solid lines represent the transient noise (*CV* ^2^(*t*)) in GFP intensity fit from the **eq. 10**, with mean decay rate (*m*_*d*_) for DMSO as 0.093 *min*^*−*1^, 0.058 *min*^*−*1^ for 1*µ*M and 0.039 *min*^*−*1^ for 2.5*µ*M of MG132 treatment. The noise in decay rates (*CV*_*d*_) were 0.25, 0.41, 0.58 for DMSO control, 1*µ*M, and 2.5*µ*M treatment of MG132, respectively.

**Figure S4.**
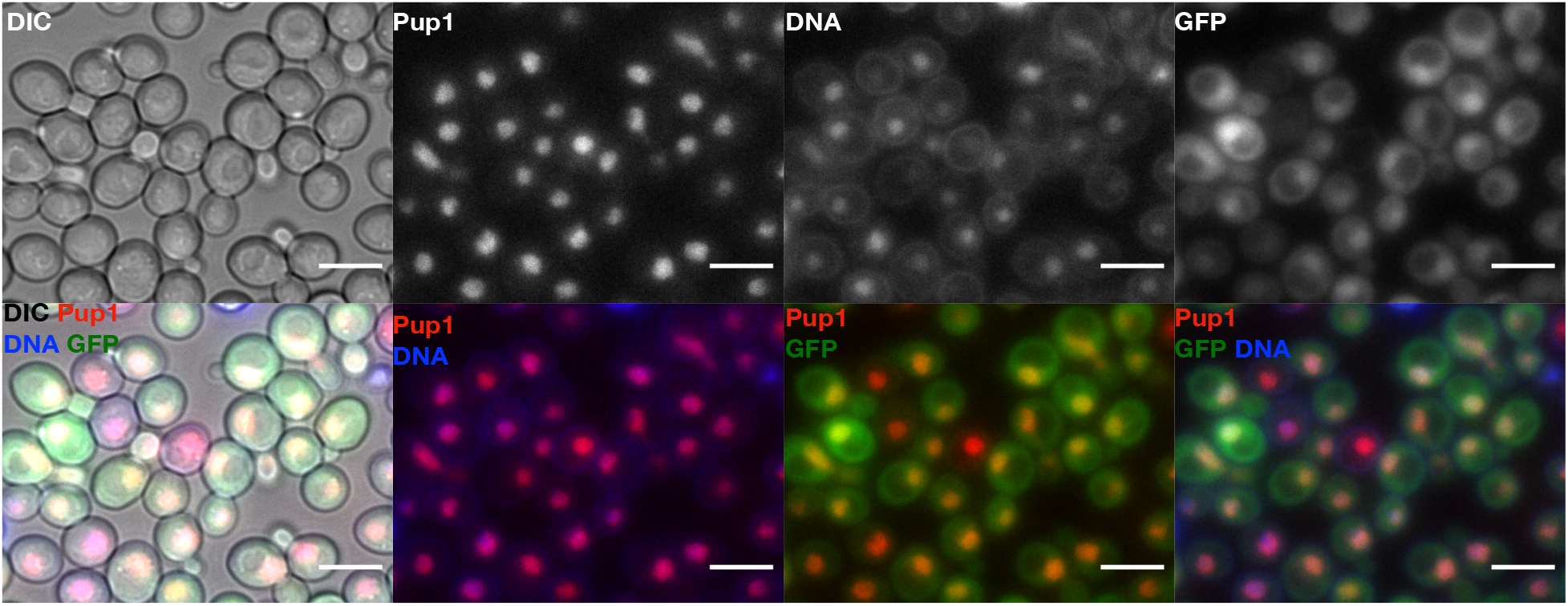
Microscopy images showing the expression of the catalytic proteasomal subunit, Pup1, tagged with a red fluorescent tag tDimer-RFP in yeGFP-mODC expressing cells. Pup1 is localized to the cell’s nucleus, as seen by colocalization with the DNA stain. The scale bar represents 6 *µm*.

**Figure S5.**
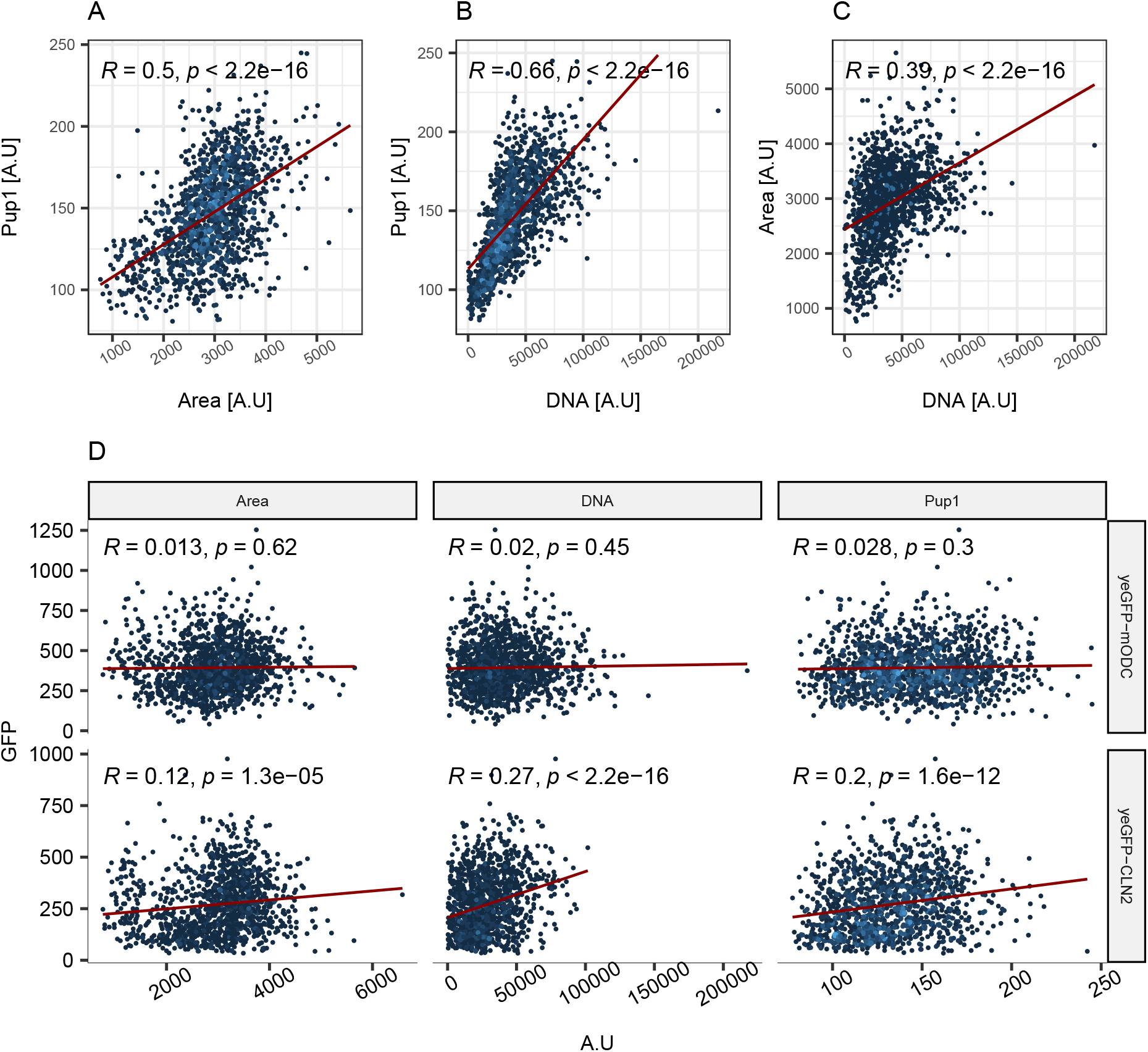
Colinearity of the cellular features. **A-C**: The amount of Pup1 in a cell scales with cell size and the DNA content of the cell. The DNA content of the cell and the cell size also scale with each other. The data is shown for yeGFP-mODC degron expressing cells. **D**: The relationship of GFP expression and cellular features for the two degron GFP in the study.

### Supplemental methods

#### Construction of the GFP expressing plasmids

The three GFPs studied were built on the backbone of the pYG026 plasmid. pYG026 was provided by Junbiao Dai (Addgene plasmid # 65328; http://n2t.net/addgene:65328; RRID: Addgene 65328) [98]. The key features of the plasmid are a constitutively expressed mCherry cassette, the CEN/ARS region to maintain a single copy of the plasmid in yeast cells, a *URA3* and *LEU2* expression cassette for auxotrophic selection, a yeGFP CDS with an ADH1 terminator and an upstream variable region to facilitate the cloning of a promoter of choice. The GFP mRNA expression in the study is controlled by a galactose inducible promoter Gal1 and by the ADH1 terminator. The construction of each plasmid expressing yeGFP, yeGFP-mODC, and yeGFP-CLN2 are summarized in **Supplemental figure SS6**.

##### yeGFP

To clone in the galactose promoter to control the expression of yeGFP, the pYG506 plasmid region was amplified with primers SD1 M and SD5 **SS6A**. The galactose promoter sequence, along with the yeGFP-mODC CDS, was commercially synthesized on a bacterial plasmid pSD001 from twist bioscience. The galactose promoter sequence was amplified from the pSD01 plasmid using primers SD6 and SD7. The PCR products of SD M + SD5 and SD6 + SD7 were used to create the plasmid pSD02 using Gibson assembly resulting in the expression of yeGFP under the control of galactose promoter.

##### yeGFP-mODC

The pSD02 plasmid was used to swap out the yeGFP CDS with yeGFP-mODC CDS **SS6B**. Briefly, the plasmid backbone was amplified with SD2 M and SD7. The yeGFP-mODC CDS was amplified from the pSD01 plasmid using SD16 and SD8 primers. The *ADH1* terminator sequence was amplified from pYG026 plasmid using SD3 and SD4 primers. The three PCR products were Gibson assembled to create the plasmid pSD03.

##### yeGFP-CLN2

Similarly, the yeGFP CDS was swapped from the pSD02 plasmid with yeGFP-CLN2 CDS from the plasmid pITGFP89 **SS6C**. pIITGFP89 was provided by Claudia Vickers (Addgene plasmid #83560 ; http://n2t.net/addgene:83560 ; RRID:Addgene 83560) [61]. yeGFP-CLN2 CDS was amplified from the pITGFP89 plasmid using SD9 and SD10 primers. The PCR products of SD2 M + SD7, SD3 + SD4, and SD9 + SD10 were Gibson assembled to create pSD04 plasmid.

#### Genomic integration

To genomically integrate the plasmids carrying the GFP expression cassette, the CEN/ARS sequence must be deleted from the pSD02, pSD03, and pSD04 plasmids. This was achieved by digesting the plasmids with PfoI and PmlI and ligating the blunt ends, **Supp. Fig. S6D**. The removal of the CEN/ARS was confirmed by sequencing with SD23 and SD20 to read into the *LEU2* and AmpR regions, respectively. To use these constructs in yeast strains with a Pup1-tDimer background, the mCherry cassette needed to be deleted from the constructs. This was achieved by the digestion of the plasmids with XbaI + BsmI, blunt end ligation of the resulting digested plasmid, and then confirming the lack of mCherry cassette using sangar sequencing, **Supp. Fig. S6D**.

The resulting plasmids pSD08, pSD09, pSD011 linearized with BstEI. The linearized product was transformed in the ySD011 strain for genomic integration of the GFP cassettes in the *LEU2* locus, and the positive transformants were selected for on the SD-ura3-leu2 selection plates. The single colonies were screened for single vs. multiple events of homologous recombination via PCR amplification from SD25 and SD24 (for a single copy of the GFP cassette), and SD23 and SD25 (for multiple copies of the GFP cassettes), **Supp. Fig. S6E**.

**Figure S6.**
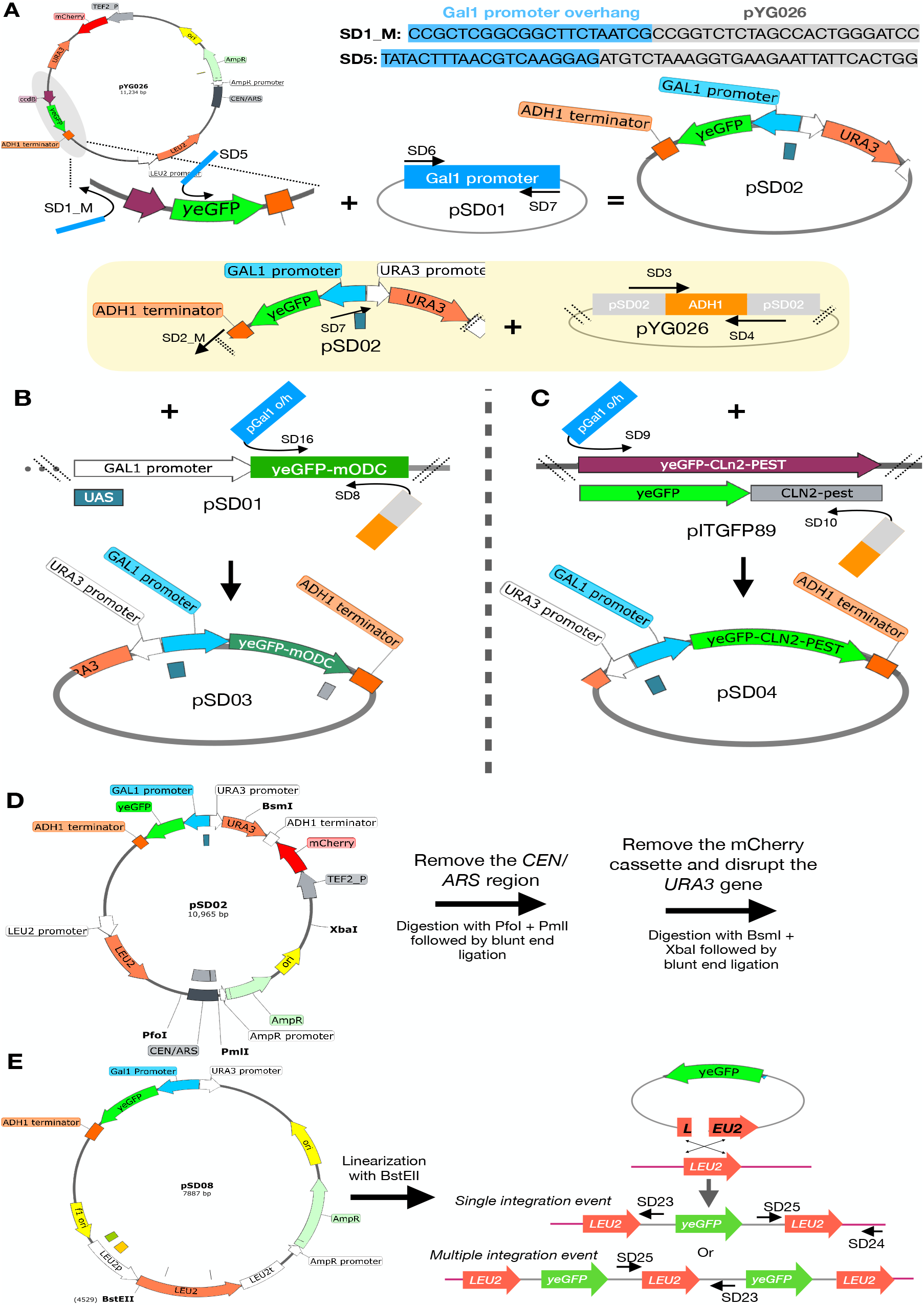
Construction of plasmids used to integrate the GFP expression cassette genomically. **A**:Construction of the pSD02 plasmid. This plasmid carries the yeGFP CDS under the control of the *GAL1* promoter and is terminated by *ADH1* terminator. pYG026 plasmid was used as the backbone. A simplified map of the plasmid is shown with the key features. The backbone of the plasmid was amplified using primers SD1 M and SD5. These primers had a 5’ overhang of GAL1 promoter sequence to facilitate Gibson assembly with the Gal1 promoter sequence amplified from the pSD01 plasmid. **B**: Construction of the pSD03 plasmid carrrying the yeGFP-mODC expression cassette. pSD02 plasmid was used as the backbone. The plasmid sequence including the *GAL1* promoter sequence was amplified using SD2 M and SD7 primers. The *ADH1* terminator sequence was excluded because there were multiple sites of annealing for the SD3 primer which when used with SD7 primer would result in the amplification of a small portion of the plasmid instead of the desired longer product. The yeGFP-mODC coding sequence was amplified from the pSD01 plasmid using primers SD16 and SD8. These primers contained 5’ overhangs to facilitate gibson assembly of the three PCR products to form the plasmid pSD03. **C**: Construction of plasmid pSD04 carrying the yeGFP-CLN2 expression cassette. The yeGFP-CLN2 CDS was amplified from the pITGFP89 plasmid using the primers SD9 and SD10. The PCR product was Gibson assembled with the amplified plasmid backbone and the *ADH1* amplified sequence. The cartoon exhibits simplified maps of the plasmid to highlight the various steps involved. **D**,**E**: Genomic integration of the GFP expression cassette. The resulting plasmids maintain the other features of the backbone pYG026 plasmid. The CEN/ARS sequence was removed via restriction digestion with PfoI and PmlI to improve the efficiency of genomic integration of the plasmid. The mcherry and the *URA3* expression cassettes were removed via restriction digestion with XbaI and BsmI. **E**: The plasmids were linearized in the *LEU2* region using BstEII for the genomic integration of the GFP expression cassette in the endogenous *LEU2* locus. The linearized plasmid was transformed in the SDY11 strain. Colonies with Single integration events were selected via a positive PCR screening using primers SD24 and SD25 and discarding colonies with a positive PCR with SD25 and SD23. The schematic for the PCR screen was adapted from [94].

**Table1:**
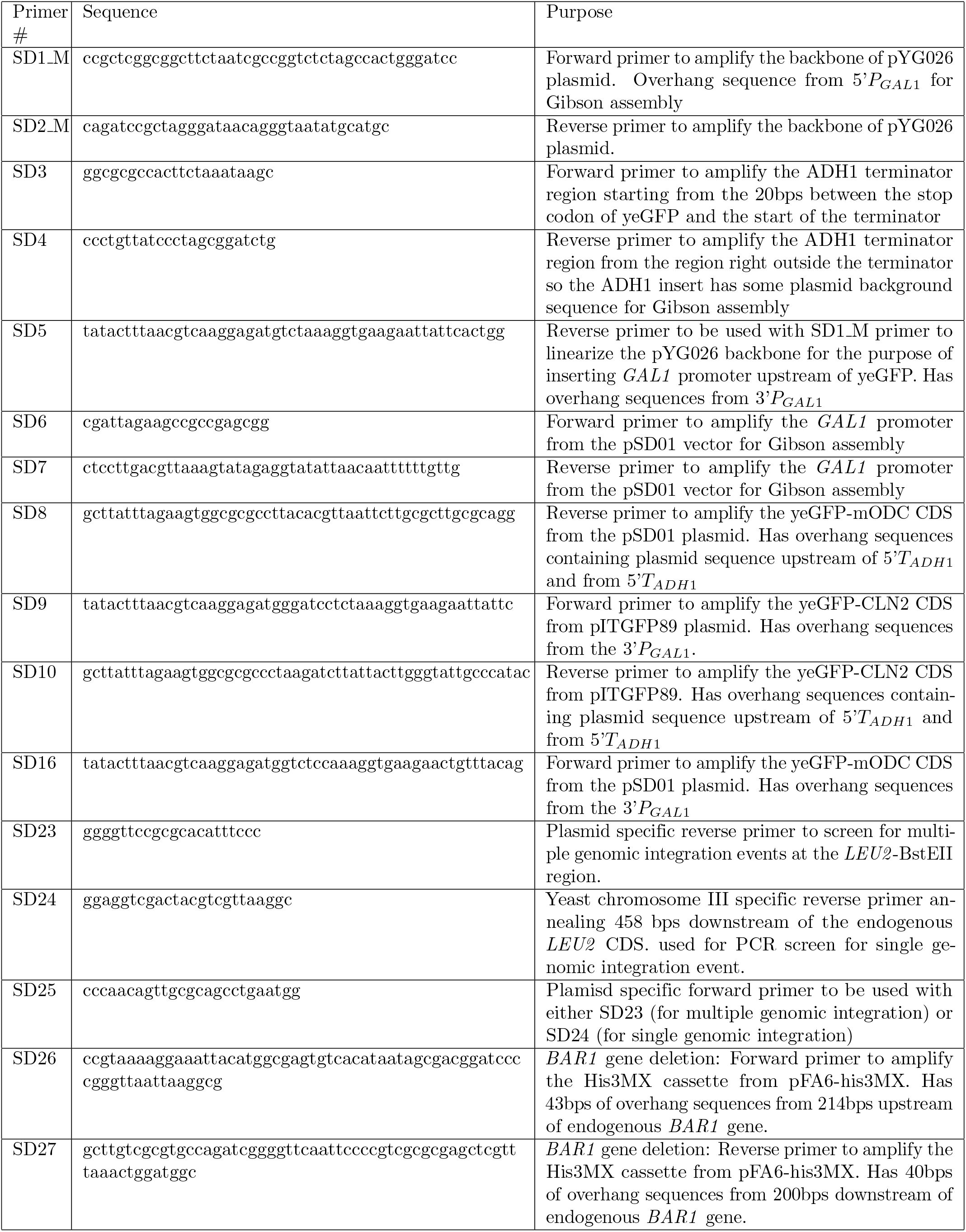
List of primers. List of primers used in the study.

**Table 2:**
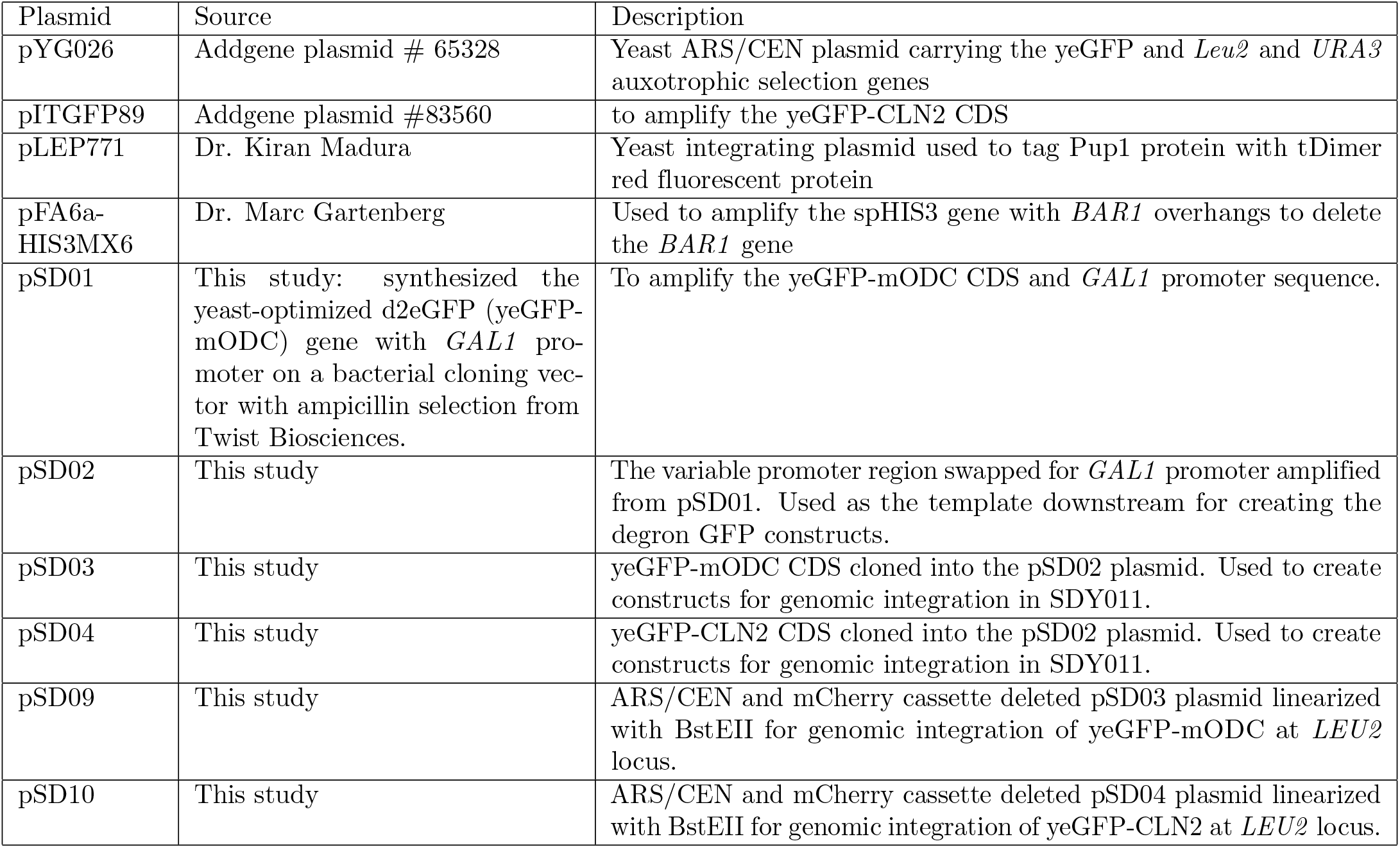
List of plasmids. List of plasmids used and generated in the study

**Table 3:**
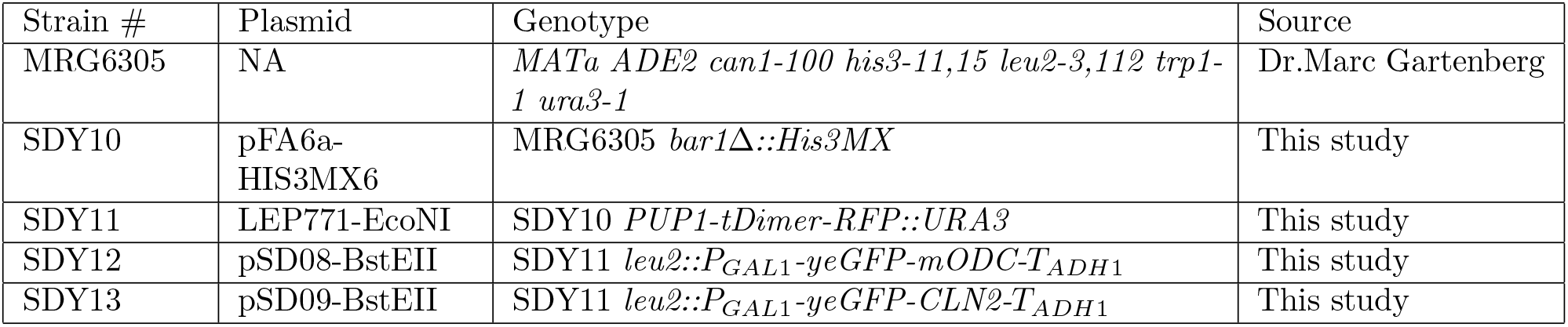
List of yeast strains. List of yeast strains used and generated in the study.

| Strain # | Plasmid | Genotype | Source |
| --- | --- | --- | --- |
| MRG6305 | NA | *MATa ADE2 can1-100 his3-11,15 leu2-3,112 trp1-1 ura3-1* | Dr.Marc Gartenberg |
| SDY10 | pFA6a-HIS3MX6 | MRG6305 *bar1Δ::His3MX* | This study |
| SDY11 | LEP771-EcoNI | SDY10 *PUP1-tDimer-RFP::URA3* | This study |
| SDY12 | pSD08-BstEII | SDY11 *leu2::P<sub>GAL1</sub>-yeGFP-mODC-T<sub>ADH1</sub>* | This study |
| SDY13 | pSD09-BstEII | SDY11 *leu2::P<sub>GAL1</sub>-yeGFP-CLN2-T<sub>ADH1</sub>* | This study |

## Supplemental files

Supplemental tables 4 and 5 were used to generate figures 2, 3, and 5, and Supplemental figures S2 and S5. Supplemental tables 6 and 7 were used to generate Figure 4 and Supplemental Figure S3. All the data presented in this study is attached as CVS files and also available in the GitHub repository: https://github.com/shahlab/ProteinDecayNoise-paper.git

## Data availability

The data used to generate Figures 2, 3, and 5 and supplemental Figures S2 and S5 are in Supplemental Tables 4 and 5. Supplemental Table 4 has the single-cell time-lapse data, and Supplemental Table 5 has the single-cell parameters of the mechanistic decay model, along with cellular attributes at *t* = 0. The data used to generate Figure 4 and the supplemental Figure S3 are in Supplemental Tables 6 and 7. Supplemental Table 6 contains the timelapse data for the protein inhibition experiment, and Supplemental Table 7 contains the estimated parameters of the decay model, along with the cellular attributes for the protein inhibition experiment. The code to generate the figures is available at GitHub https://github.com/shahlab/ProteinDecayNoise-paper.git. The raw and processed images are uploaded to Dryad (DOI):doi:10.5061/dryad.bnzs7h4g6.

## Acknowledgments

A.S. acknowledges support from NIGMS (NIH) under award number R35GM148351. S.D. and P.S. were supported by NIH/NIGMS grant R35 GM124976 and start-up funds from the Human Genetics Institute of New Jersey at Rutgers University.

## Competing interests

Premal Shah receives compensation, holds equity, and is a Director at Ananke Therapeutics. S.D. and A.S. declare no competing interests.

